# Mapping genetic effects on cell type-specific chromatin accessibility and annotating complex trait variants using single nucleus ATAC-seq

**DOI:** 10.1101/2020.12.03.387894

**Authors:** Paola Benaglio, Jacklyn Newsome, Jee Yun Han, Joshua Chiou, Anthony Aylward, Sierra Corban, Mei-Lin Okino, Jaspreet Kaur, David U Gorkin, Kyle J Gaulton

## Abstract

Gene regulation is highly cell type-specific and understanding the function of non-coding genetic variants associated with complex traits requires molecular phenotyping at cell type resolution. In this study we performed single nucleus ATAC-seq (snATAC-seq) and genotyping in peripheral blood mononuclear cells from 10 individuals. Clustering chromatin accessibility profiles of 66,843 total nuclei identified 14 immune cell types and sub-types. We mapped chromatin accessibility QTLs (caQTLs) in each immune cell type and sub-type which identified 6,248 total caQTLs, including those obscured from assays of bulk tissue such as with divergent effects on different cell types. For 3,379 caQTLs we further annotated putative target genes of variant activity using single cell co-accessibility, and caQTL variants were significantly correlated with the accessibility level of linked gene promoters. We fine-mapped loci associated with 16 complex immune traits and identified immune cell caQTLs at 517 candidate causal variants, including those with cell type-specific effects. At the 6q15 locus associated with type 1 diabetes, in line with previous reports, variant rs72928038 was a naïve CD4+ T cell caQTL linked to *BACH2* and we validated the allelic effects of this variant on regulatory activity in Jurkat T cells. These results highlight the utility of snATAC-seq for mapping genetic effects on accessible chromatin in specific cell types and provide a resource for annotating complex immune trait loci.

## Introduction

Genome-wide association studies have identified thousands of genomic loci associated with complex human traits and disease^1–3^, but their molecular mechanisms remain largely unknown. Interpreting the mechanisms of trait-associated loci is paramount to an improved understanding of the cell types, genes and pathways involved in complex traits and disease^1^. Genetic variants at complex trait-associated loci are primarily non-coding and enriched in transcriptional regulatory elements^1,4,5^, implying that the majority affect gene regulatory programs. As gene regulation is highly cell type-specific^6,7^, uncovering the molecular mechanisms of complex trait loci requires determining the function of non-coding variants in the individual cell types that comprise a tissue. While substantial advances have been made in annotating the non-coding genome^5,8^, the regulatory effects of genetic variants in specific cell types are still largely unknown.

Mapping quantitative trait loci (QTLs) for molecular phenotypes such as gene expression levels, histone modifications and chromatin accessibility is an effective strategy to determine the regulatory activity of genetic variants^9–15^. Molecular QTL studies to date have been primarily performed in ‘bulk’ tissue, cell lines, or individual sorted cell types, however, and therefore have not yet widely annotated the breadth of cell type effects. Single cell technologies have created new avenues to study gene regulation in the specific cell types comprising a heterogeneous tissue and define relationships to complex traits and disease^16,17^. Several recent studies mapped gene expression QTLs (eQTLs) using cell type-specific expression profiles derived from single cell RNA-seq assays^18–20^. These studies represented proof-of-concept for using profiles derived from single cell data to map genetic effects on molecular phenotypes in specific cell types and sub-types. Moreover, they enabled additional analyses which leveraged data from across thousands of cells such as the identification of co-expression QTLs^19^. To date, however, no studies have mapped chromatin accessibility QTLs in specific cell types and sub-types using single cell assays.

In this study we used single nucleus ATAC-seq (snATAC-seq) to profile human peripheral blood mononuclear cell (PBMC) samples. We derived chromatin accessibility profiles of immune cell types and sub-types and mapped chromatin accessibility QTLs (caQTLs) for these profiles which identified thousands of immune cell type and sub-type caQTLs. We characterized caQTLs for each cell type, including caQTLs whose effects are obscured in bulk assays, and linked distal caQTLs to putative target gene promoters using single cell co-accessibility. Finally, we fine-mapped causal variants at genomic loci associated with 16 complex immune traits and diseases, annotated fine-mapped variants for these traits with immune cell type caQTLs and validated the molecular effects of high-probability caQTL variants.

## Results

### Chromatin accessibility profiling of peripheral blood mononuclear cells

We performed snATAC-seq and genotyping of human peripheral blood cell (PBMC) samples in order to map genetic effects on lymphoid and myeloid cell type accessible chromatin (**Figure 1a**). We used droplet-based snATAC-seq (10X Genomics) to assay 10 PBMC samples from individuals of self-reported European descent (**Supplementary Table 1, see Methods**). The snATAC-seq libraries were sequenced to an average depth of 178M reads, and libraries had consistently high-quality metrics including enrichment at transcription start sites (TSS) and fraction of reads mapping in peaks (**Supplementary Table 2**). We then performed array genotyping of each sample and imputed genotypes into 39.6M variants in the Haplotype Reference Consortium (HRC) panel^21^. Principal components analysis of genotypes mapped onto 1000 Genomes Project data confirmed European ancestry for the majority of samples (**Supplementary Figure 1**).

**Figure 1.**
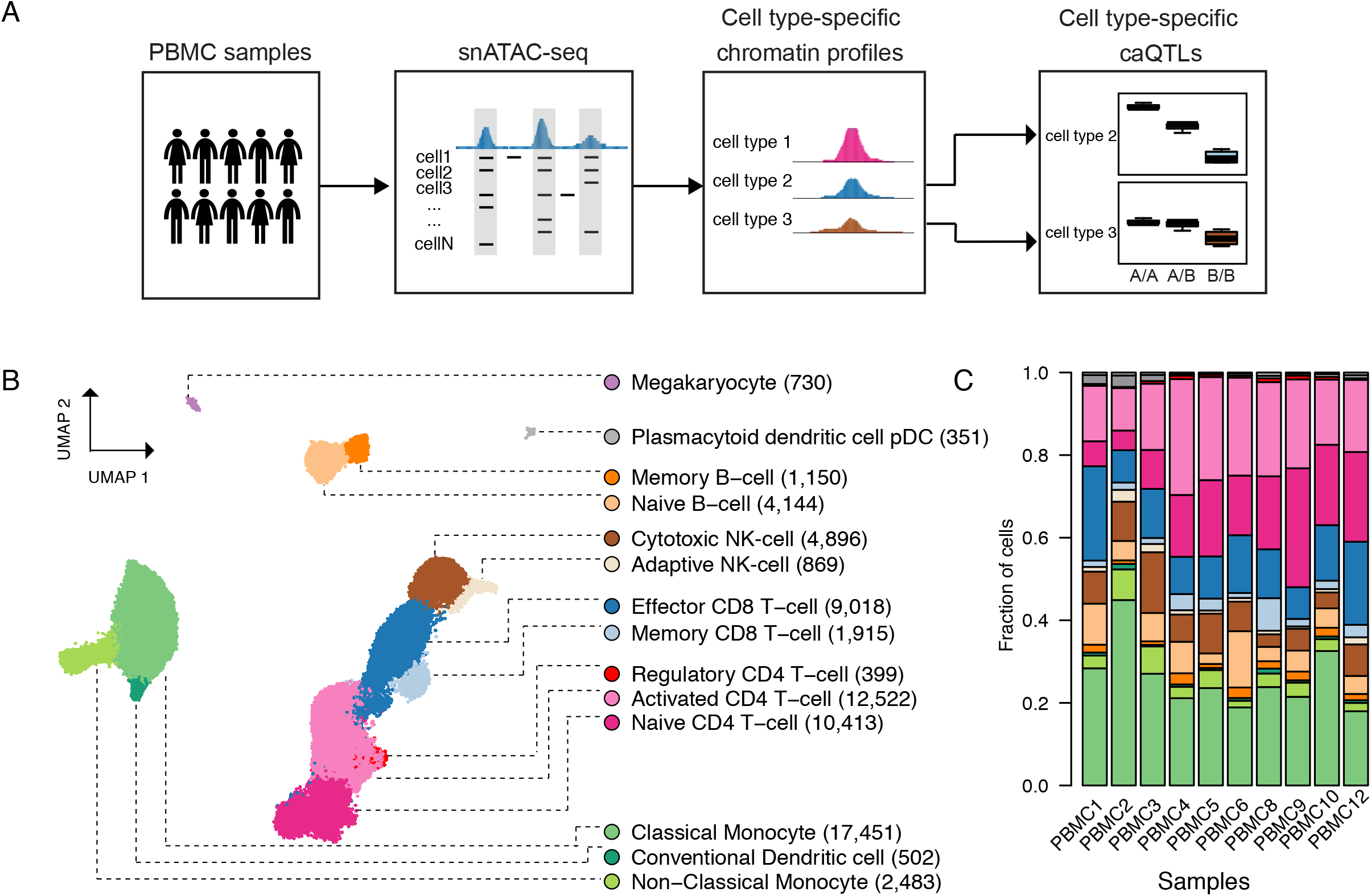
Single nucleus ATAC-seq in a population of PBMC samples. a) Schematic overview of the study. b) Clustering of single cell accessible chromatin profiles of 66,843 PBMCs from 10 individuals. Cells are plotted based on the first two UMAP components. Fourteen distinct clusters, indicated by different colors, were identified and assigned to a cell type based on known marker genes. The number of cells for each cell type is indicated in parenthesis. c) Barplot showing the relative proportions of each cell type in each sample. Color scheme is the same as in 1b.

After extensive quality control that removed low quality cells and potential doublet cells (**see Methods**), we performed clustering of 66,843 snATAC-seq profiles, which revealed 14 clusters (**Figure 1b**). We then assigned clusters to lymphoid and myeloid cell types and sub-types based on the chromatin accessibility patterns at known marker genes (**Figure 1b, Supplementary Figure 2, Supplementary Table 3**). For example, among immune cell types, *NCR1* accessibility marked NK cells, *MS4A1* accessibility marked B cells, and *PCTRA* accessibility marked plasmacytoid dendritic cells. Among cell sub-types, accessibility at *FOXP3* differentiated regulatory T cells from other T cell sub-types, and accessibility at *TCL1A* differentiated naïve B cells from memory B cells (**Supplementary Figure 2**). The proportion of each immune cell type and sub-type was broadly consistent across samples (**Figure 1c, Supplementary Figure 3a**) and was highly correlated with cell proportions determined from flow cytometry of cell surface markers for each sample (**Supplementary Figure 3b-c**). Similarly, clusters were composed of similar proportions of cells from different individuals (**Supplementary Figure 3d**). These results demonstrate that snATAC of PBMCs resolved lymphoid and myeloid cell types and sub-types with broadly consistent representation across samples.

### Mapping chromatin accessibility QTLs in immune cell types and sub-types

Within each immune cell type and sub-type cluster, we aggregated reads for all cells in the cluster, generated accessible chromatin read count profiles, and called accessible chromatin sites using MACS2^22^. Considering all immune cell types and sub-types there were 210,771 total accessible chromatin sites (**Supplementary Table 5**). Immune cell type and sub-type sites were highly concordant with sites identified in a previous study of FACS-sorted immune cell types^23^ (**Supplementary Figure 4**). We then performed QTL mapping of chromatin accessibility read counts in these sites using RASQUAL^24^, a method which combines population-based and allele-specific mapping. We focused on the 5 immune cell types with appreciable numbers of cells (B, CD4+ T, CD8+ T, monocyte, NK) and mapped QTLs at both cell type and sub-type resolution. For each cell type or sub-type, we retained sites with >5 reads per sample on average and only tested variants that mapped directly in accessible sites and were heterozygous in at least two samples. After applying these filters, on average 67,979 variants per cell type were tested for association with 46,373 peaks (3.8 variants/peak). For comparison, we also performed caQTL mapping after merging all reads for each sample ignoring their cell of origin to mimic a ‘bulk’ ATAC-seq experiment.

In total we identified 6,248 distinct caQTLs in an immune cell type or sub-type (at FDR <0.1), including 5,187 at cell type resolution and 5,398 at sub-type resolution (**Figure 2a, Supplementary Table 6**). We also identified 5,697 caQTLs at ‘bulk’ resolution (**Figure 2a, Supplementary Table 6**). There was limited evidence for reference bias in the resulting caQTLs (1.68% with ψ<.25, annotated in **Supplementary Table 6**). Excluding the allelic imbalance component from QTL mapping resulted in substantially fewer caQTLs at FDR<.10 (168 cell type, 426 sub-type) although the allelic effects were highly concordant (**Supplementary Figure 5**). The majority of caQTLs were identified at FDR<.10 at different resolutions, although a subset was found only at one resolution (**Figure 2a**). The number of caQTLs identified in each cell type was proportional to the number of cells for that cell type (**Figure 2b**), likely due to differences in available read depth leading to reduced power for less common cell types. Most caQTLs were identified at FDR<.10 in only one cell type or sub-type (80% for cell types, 75% for cell sub-types) **(Figure 2c**). However, when considering caQTLs significant in at least one cell type, allelic effects (π) were strongly correlated across cell types as well as with ‘bulk’ data (median Spearman correlation *r*=0.63, **Figure 2d**), with stronger correlation between more similar cell types.

**Figure 2.**
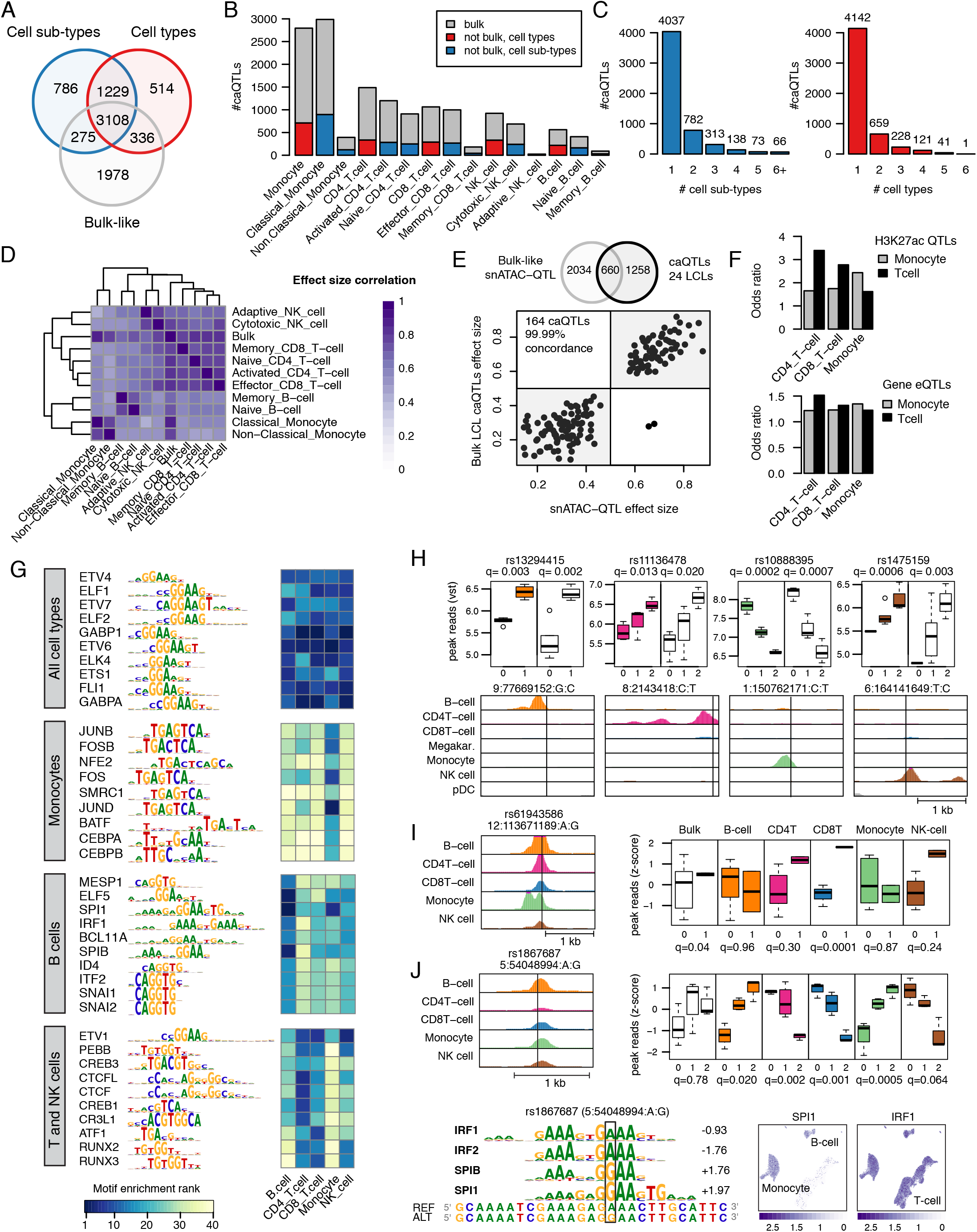
Identification and characterization of immune cell type chromatin accessibility QTLs. a) Venn diagram showing the total number of caQTLs from single-cell ATAC-seq across immune cell types (red), cell sub-types (blue); and in ‘bulk’ (gray). b) Number of caQTLs identified in each cell type (red for cell types, blue for sub-types) and the subset found in ‘bulk’ data (gray). c) Number of caQTLs unique or common to different cell types. d) Heatmap of pairwise correlation (Spearman) between effect sizes of caQTLs, where association is significant in at least one of the two cell types in the pair. e) Comparison between caQTLs from PBMC bulk-like data and published caQTLs from 24 LCLs. Venn diagram on top indicates the number of significant caQTLs in each dataset and their overlap. Scatter plot of effect sizes for caQTLs found in both studies and having the same lead variant. f) Overlap between cell type caQTLs and H3K27ac QTLs (top) or gene expression QTLs (bottom) in either Monocytes (gray) or T cells (black). g) Top transcription factor motifs disrupted by caQTL variants across different cell types. Clustering is based on motif similarity. The heatmap shows the enrichment ranking of each TFs in each cell type. h) Examples of caQTLs in peaks specific to a single cell type including rs13294415 (B cell-specific), rs1475159 (CD4+ T cell-specific), rs10888395 (Monocyte-specific) and rs11136478 (NK cell-specific). Top panels: colored-coded box-plots show association in the different cell types, white box-plots show caQTL in ‘bulk’ PBMCs. Association q-values are shown on the top and variant genomic location (hg19) is shown at the bottom. Bottom panels: genome-browser screenshot of cell type chromatin profiles. i) Variant rs61943586 was in peaks active in all cell types but was a significant caQTL in CD8+ T cells only. Left: genome-browser screenshot of cell type chromatin signal. Right: boxplots as in H. l) Variant rs1867687 was a significant caQTL in all cell types but had opposite effects in different cell types. Top-left: genome-browser screenshot of cell type chromatin signal. Top-right: boxplot of signal split by genotype in bulk and each cell type. Boxplots are color-coded as in I. Bottom-left: TF motifs altered by the rs1867687 variant and their respective score differences are shown. Positive scores indicate preference for alternate allele. Bottom-right: UMAP plot showing accessibility of the SPI1 gene in Monocytes and B-cells and ubiquitous accessibility of IRF1.

We next compared cell type caQTLs in our study to external QTL datasets previously generated in immune cells. We first compared caQTL results from our ‘bulk’ analysis with caQTLs previously mapped in 24 lymphoblastoid cell lines (LCLs)^24^. Of the 2,694 caQTLs in our data that were tested in the LCL study, 660 (24.4%) were also significant LCL caQTLs (OR=15.9, P<2.2×10^−16^, Fisher’s exact test) of which 164 shared also the same lead variant (OR=4.2, P=1.05×10^−16^ Fisher’s exact test) and were 99.99% concordant in their effect direction (**Figure 2e**). Of note, when considering caQTLs from each individual cell type, B cell caQTLs had the highest overlap with LCL caQTLs, consistent with LCLs being derived from B cells (**Supplementary Figure 6**). We next compared caQTLs in our study to published histone H3K27ac QTLs (hQTLs) and expression QTLs (eQTLs) from FACS-sorted T cells and Monocytes from the BLUEPRINT project^15^. The enrichment for T cell hQTLs was stronger in CD4+ (OR=3.8, *P*=3.2×10^−113^, Fisher’s exact test) and CD8+ (OR=3.4, *P*=1.3×10^−60^) T cell caQTLs compared to monocyte caQTLs (OR=1.6, *P*=9.1×10^−16^) (**Figure 2f**). We observed the converse pattern for monocyte hQTLs, which were more enriched for monocyte caQTLs than T cell caQTLs (CD4+ T cell OR=1.65, *P*=3.8×10^−20^; CD8+ T cell OR=1.7, *P*=1.4×10^−18^; monocyte OR=2.4, *P*=1.1×10^−153^) (**Figure 2f**). We observed the same cell type enrichment pattern for T cell and monocyte eQTLs (T-cell eQTLs: CD4+T-cell OR=1.51, *P*=1.2×10^−42^; CD8+ T cell OR=1.3, *P*=5.6×10^−14^; monocyte OR=1.2, *P*=2.0×10^−15^; monocyte eQTLs: CD4+ T cell OR=1.2, *P*=2.4×10^−10^; CD8+ T cell OR=1.2, *P*=1.4×10^−8^; monocyte OR=1.3, *P*=3.9×10^−39^; **Figure 2f**).

To identify transcription factors (TFs) mediating immune cell type caQTLs, we identified TF sequence motifs preferentially disrupted by caQTL variants in each cell type. We used MotifBreakR^25^ to predicted allelic effects of SNPs on TF motifs from the HOCOMOCO v10 human database^26^, comprising 640 motifs corresponding to 595 unique TFs. We first predicted allelic motif effects for all variants tested for QTL association. Then, for each TF motif, we compared the proportion of motif instances disrupted by caQTLs compared to non-caQTL variants. Thus, we were able to measure the enrichment of predicted TF-disrupting caQTLs for each TF motif. Immune cell type caQTLs were broadly enriched for disrupting any TF motif compared to non-caQTL variants (OR=1.2, *P*=6.1×10^−4^, Fisher’s exact test). When considering caQTLs in each cell type, there were 25 TF motifs significantly enriched for B cell caQTLs, 44 motifs enriched for CD4+ T cell caQTLs, 29 motifs enriched for CD8+ T cell QTLs, 29 motifs enriched for NK cell QTLs and 93 motifs enriched for monocyte QTLs (FDR<0.05, one-tailed binomial test, **Figure 2g, Supplementary Table 7**). Motifs disrupted by caQTLs included those with broadly shared enrichment across different cell types including ETS1, ETV6 and GABP1, as well as those with highly cell type-specific enrichment such as BCL11A in B cells (FDR= 0.012), SPI1 in B cells and monocytes (FDR=3.07×10^−8^, FDR=5.10×10^−34^), and CEBPB in monocytes (FDR=7.8×10^−17^).

At numerous loci, caQTLs mapped at cell type and sub-type resolution provided insight beyond those obtained by mapping caQTLs in bulk tissue. The most straightforward examples consisted of caQTLs for accessible chromatin sites active in only one cell type, where the effects of a caQTL identified in bulk data could be simply ascribed to that cell type (1,776 caQTLs, **Supplementary Figure 7a**). For example, rs13294415 was a caQTL for a B cell-specific site (allelic effects [π]=.79, q-value=.003), rs11136478 was a caQTL for a CD4+ T cell-specific site (π=.65, q=.013), rs10888395 was a caQTL for a monocyte-specific site (π=.31, q=2.1×10^−4^) and rs1475159 was a caQTL for a NK cell-specific site (π=.81, q=6.3×10^−4^) (**Figure 2h**). We also identified caQTLs for immune sub-type-specific sites (1,325 caQTLs), such as rs3014874 which was a caQTL for a classical monocyte-specific site (π=.28, q=.008) and rs7094953 which was a caQTL for a naïve CD4+ T cell-specific site (π=.26, q=.008) (**Supplementary Figure 7b**). Another class of caQTLs were those for sites active in all cell types, yet where the variant effects were specific to only a few cell types (2,362 and 2,704 for cell types and subtypes, respectively). For example, variant rs61943586 mapped in a site active in all immune cell types and had a significant effect in CD8+ T cells (π=.76, q=1.4×10^−4^), but no effect in B cells and monocytes (π=.50, q=.96; π=.48, q=.76) (**Figure 2i**). Similarly, variant rs747748 mapped in a site active in all cell types yet only had a significant effect in classical monocytes (π=.43, q=.0018) (**Supplementary Figure 7c**). In these latter examples, variant effects in bulk data were dampened due to the inclusion of cell types with no effect (rs61943586 bulk π=.57, q=.04; rs747748 bulk π=.48, q=.18) (**Figure 2i, Supplementary Figure 7c)**.

We also observed caQTLs with more complex effects, such as those with divergent effects on different cell types (41 and 60 for cell types and subtypes, respectively). In one example, variant rs1867687 was a significant caQTL in all immune cell types, where the G allele had increased accessibility in B cells and monocytes (B cell π=.66, q-value=.02; monocyte π=.66, q=4.8×10^−4^) and the A allele had increased accessibility in CD4+ and CD8+ T cells and NK cells (CD4+ T π=.29, q=.002; CD8+ T π=.20, q=.001; NK cell π=.27, q=.064) (**Figure 2j**). In comparison, rs1867687 had no effect in ‘bulk’ data (π=.51, q=.78). The alleles of this variant were predicted to bind different TFs, where the G was predicted to bind SPI1 and SPIB motifs and the A allele was predicted to bind IRF TF motifs (**Figure 2k**). SPI1 and SPIB motifs were specifically enriched in B cells and monocytes, whereas IRF motifs were broadly enriched across cell types (**Figure 2k**), suggesting a potential mechanism through which this variant has opposing effects on different immune cell types.

### Linking distal caQTLs to effects on target gene promoters

Among the 6,248 caQTLs identified in our study, a minority (17%) mapped to gene promoter regions. The remaining caQTLs were in chromatin sites distal to promoters, and we therefore sought to define the target genes of these caQTLs. Co-accessibility between pairs of accessible chromatin sites across single cells has been used to annotate putative target genes of distal enhancers^17,27^. We therefore defined co-accessible sites (co-accessibility score >.05) in the 5 immune cell types with >1k cells (CD4+ T, CD8+ T, B, monocyte, NK) using Cicero^27^. For each cell type we retained co-accessible sites greater than 10kb apart and that also were co-accessible in at least two samples individually. In total we identified 481,963 pairs of co-accessible sites, which included between 75k and 132k per cell type (**Figure 3a**). We compared co-accessible sites for each cell type to chromatin interactions from promoter capture Hi-C (pCHi-C) data previously generated in 16 immune cell types and sub-types^28^. We observed strongest enrichment of cell type co-accessible sites for the corresponding cell type in pCHi-C interactions in each case, except for NK cells, which were not assayed by pCHi-C (**Figure 3b**). When segregating co-accessible sites by distance, there remained strong enrichment for pCHi-C interactions even at distances of up to 1MB (**Figure 3c**).

**Figure 3.**
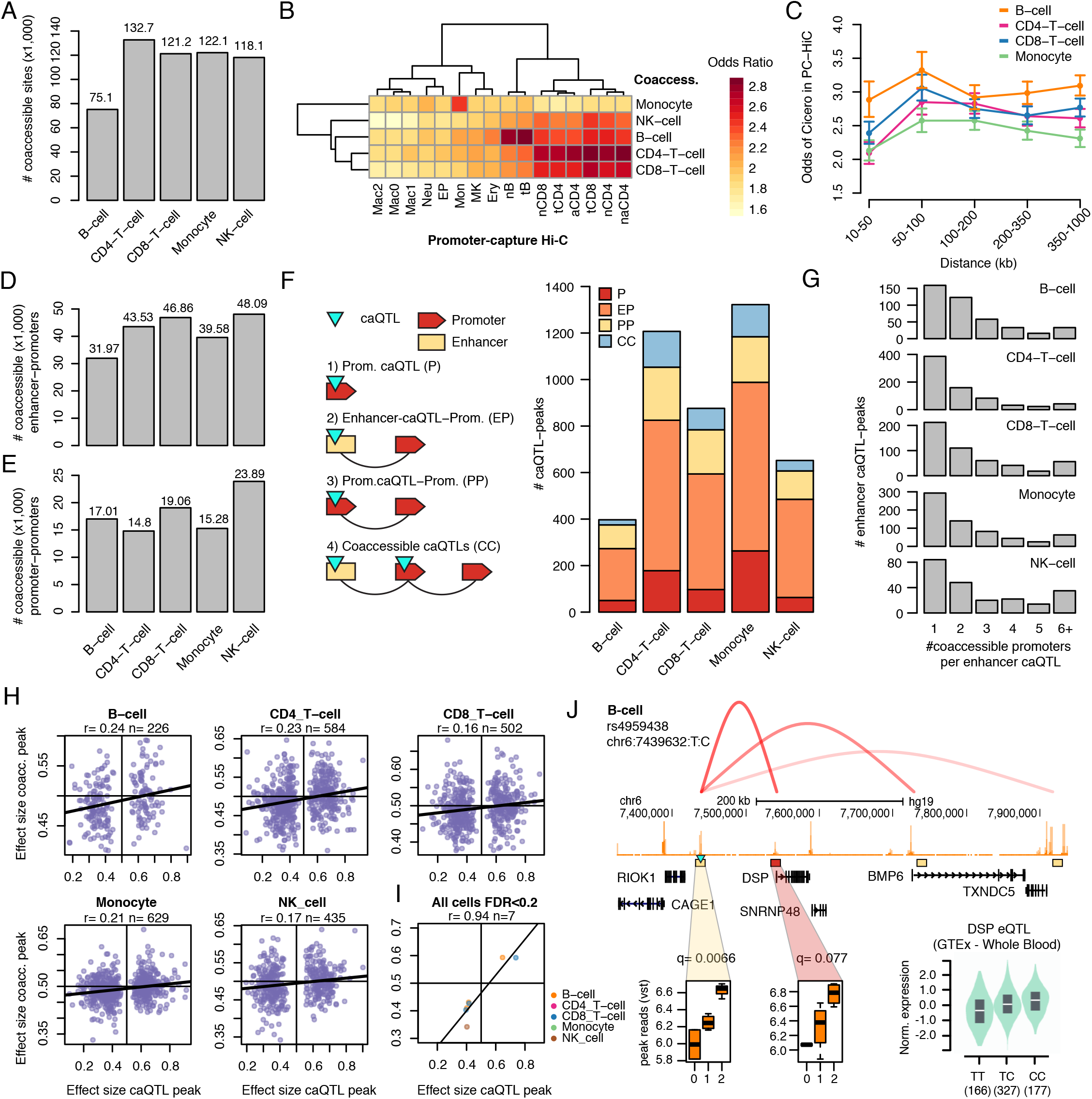
Linking distal immune cell caQTLs to putative target genes. a) Number of co-accessible links in each immune cell type. b) Enrichment of cell type co-accessible links for overlap with promoter-capture Hi-C (pcHi-C) interactions in immune cell types. c) Enrichment of cell type co-accessible links for pcHi-C interactions separated by distance between linked sites. d) Number of co-accessible links between a promoter site (+/- 1kb) and a distal (non-promoter) site in each cell type. e) Number of co-accessible links between promoter sites. f) Breakdown of caQTLs linked to promoters in each cell type, including caQTLs directly in promoter sites, caQTLs in distal sites co-accessible with a promoter site, caQTLs in promoter sites co-accessible with a different promoter site, and more complex cases involving multiple linked caQTLs. g) Breakdown of caQTLs in each cell type by the number of promoter sites they were linked to. h) Correlation in the effects of caQTL variants on the primary site and co-accessible promoter sites in each cell type. Pearson correlation coefficient and number of co-accessible pairs of peaks are indicated. i) Correlation in caQTL variant effects on the primary site and co-accessible promoter site for variants significant (FDR 20%) for the latter. j) A caQTL in B cells rs4959438 was also a QTL for a co-accessible site at the *DSP* promoter and an eQTL for *DSP* in GTEx.

Using the co-accessible sites identified for each cell type we then annotated caQTLs with their putative target genes. There were 179,347 distal accessible chromatin sites co-accessible with at least one promoter site (30.5k-44.4k per cell type) and 66,571 promoter sites co-accessible with promoter sites of a different gene (13.0k–18.1k per cell type) (**Figure 3d-e**). Across all 6,248 caQTLs, 3,379 were either in a site co-accessible with at least one gene promoter or in a promoter site directly. Among these 3,379 caQTLs, the majority were distal sites co-accessible with a promoter (54-65% per cell type) (**Figure 3f**. Among distal caQTLs co-accessible with a gene promoter, 38-53% were linked to just one gene (**Figure 3g**).

Previous studies have identified coordinated allelic effects between distal sites and interacting promoters^29^. We therefore tested caQTL variants for association with chromatin accessibility levels of all promoter sites co-accessible with the caQTL site. There was a positive and highly significant correlation between variant allelic effects on the original site and effects on co-accessible promoter sites (B r=.24, CD4+ T r=.23, CD8+ T r=.16, monocyte r=.21, NK r=.17, Pearson correlation) (**Figure 3h**). When separating co-accessible sites by distance, the correlations were reduced between more distal sites (**Supplementary Figure 8**). As we were unable to leverage allelic imbalance in this analysis, our power was more limited, and we only identified 7 linked promoter caQTLs at FDR<.20 (**Figure 3i**). There was a significant, positive correlation in variant effects on the linked promoter caQTL and the original caQTL (**Figure 3i**). For example, at the 6p24 locus rs4959438 was a caQTL for a distal site in B cells (π=.65, q=.0066) and was also a caQTL for the *DSP* promoter linked to the distal site (π=.59, q=.077) (**Figure 3j**). This variant was also a QTL for the expression of *DSP* in whole blood in GTEx v8^9^ (NES=.29, P=3.2×10^−12^), and which was directionally consistent with the C allele having increased activity. Together these results demonstrate how snATAC-seq data can be used to link caQTLs to effects on putative target genes.

### Identifying caQTLs at fine-mapped variants for complex immune trait loci

Genomic loci affecting complex immune traits and disease are primary non-coding, and the causal variants and molecular mechanisms at these loci are largely unknown. We therefore used immune cell type and sub-type caQTLs to annotate variants associated with complex immune traits and disease. We first collected published genome-wide association summary statistics for 16 blood cell count, autoimmune, inflammatory and allergy traits imputed into reference panels with comprehensive variant coverage such as 1000 Genomes or the Haplotype Reference Consortium (**Figure 4a, Supplementary Table 8**). At most traits, fine-mapping of causal variant sets at associated loci was either not performed as part of the initial study or not made publicly available. We therefore fine-mapped primary association signals at loci reported for these 16 traits using a Bayesian approach, from which we generated credible sets of variants representing 99% of the total posterior probability for each signal (**see Methods, Supplementary Data 1**). Across all traits there were 1,275 total credible sets, which contained a median of 16 variants, where traits with the smallest credible set sizes included monocyte count (median 6.5 variants), basophil count (median 7.5 variants) and rheumatoid arthritis (median 9 variants). At 396 signals fine-mapping resolved credible sets to 5 or fewer variants (**Figure 4a**).

**Figure 4.**
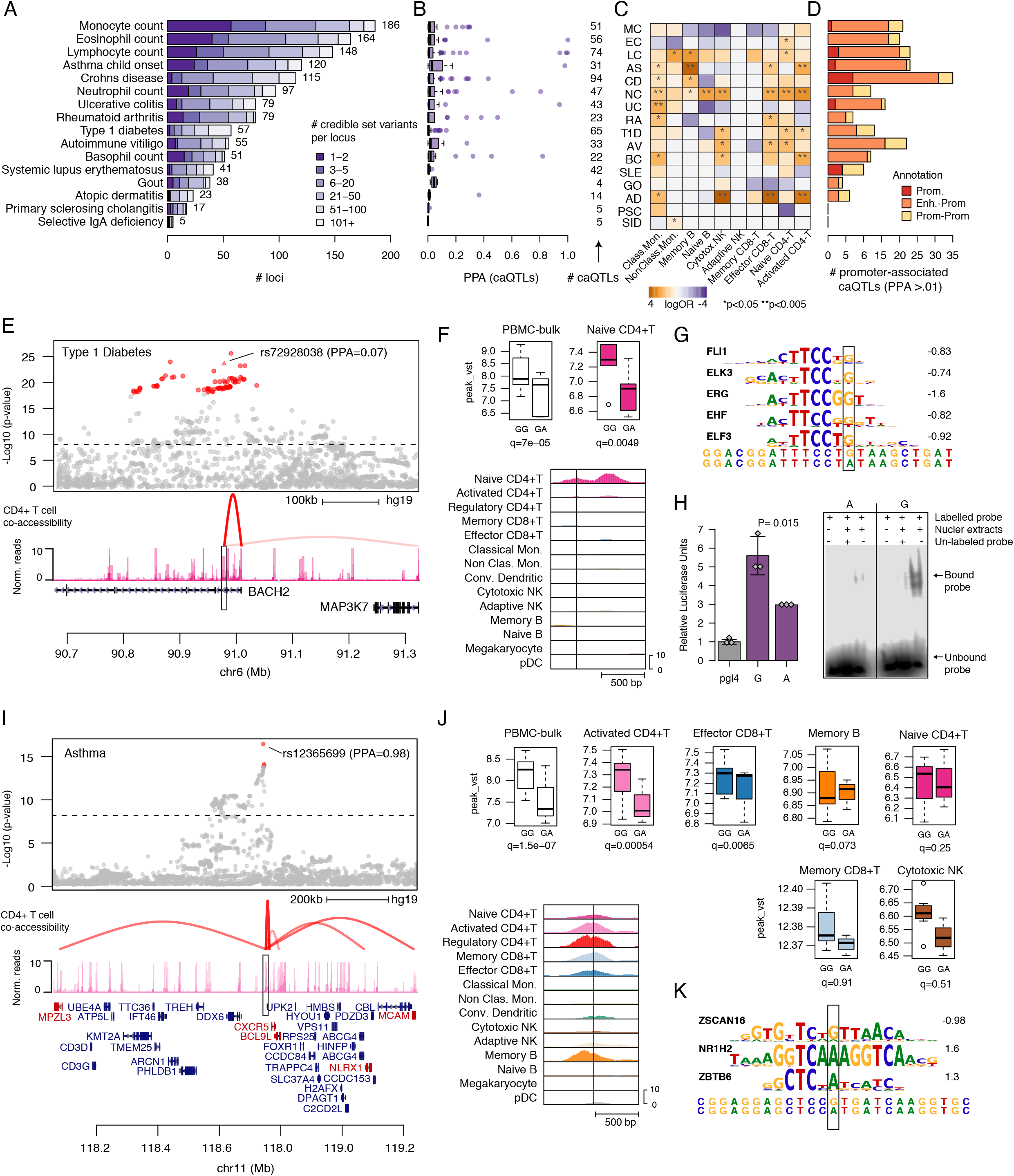
Immune cell type caQTLs at fine-mapped complex immune trait loci. a) Fine-mapping of causal variants at association signals for 16 complex immune traits and diseases. Bar plots represent the number of variants in credible sets for each trait. b) Posterior probabilities of caQTLs at fine-mapped variants for each immune trait and disease. c). d) Breakdown of caQTLs for fine-mapped variants that were linked to promoter sites. e) Regional plot of the locus on chr6 near *BACH2* associated with type 1 diabetes (T1D). (top) T1D variant association, with credible set variants highlighted in red, (bottom) chromatin signal in naïve CD4+ T cells and the co-accessible link between the site harboring rs72928038 and the *BACH2* promoter. f) Cell type-specific effects of rs72928038 on naïve CD4+ T cell chromatin. (top) Chromatin signal grouped by rs72928038 genotype in bulk PBMCs and naïve CD4+ cells on top, (bottom) genome browser of chromatin signal in each cell type. g) Predicted TF sequence motifs at rs72928038, where the variant base is highlighted. h) Validation of allelic activity for rs72928038 in T cells. (top left) Luciferase gene reporter of sequence surrounding the G and A allele of rs72928038 in Jurkat T cells. The G allele had significantly higher reporter activity. (top right) Electrophoretic mobility shift assay of oligonucleotides containing the G and A allele of rs72928038 in Jurkat T cells, where the G allele had protein binding. i) Regional plot of the chr11 locus near *CXCR5* associated child asthma. (top) Asthma association statistics, with the two credible set variants highlighted in red, (bottom) chromatin signal in activated CD4+ T cells and the co-accessible links between the site harboring rs12365699 and the *CXCR5, BCL9L, NLRX1 MCAM*, and *MPZL3* promoters. j) Cell type-specific effects of rs12365699. (top) Chromatin signal grouped by rs12365699 genotype in bulk PBMCs and activated CD4+ cells on top, (bottom) genome browser of chromatin signal in each cell type. k) Predicted TF sequence motifs at rs12365699, where the variant base is highlighted.

A total of 523 credible set variants representing 211 association signals were immune cell type or sub-type caQTLs (**Figure 4b**). We determined whether fine-mapped variants for each trait were preferentially enriched for caQTLs from specific immune cell sub-types by comparing to a background of non-caQTL sites (**see Methods**). The majority of traits (12/16) showed nominal enrichment (P<.05) for caQTL peaks in at least one immune cell sub-type, several of which recapitulated known biology of cell types contributing to the trait (**Figure 4c**). For example, type 1 diabetes (T1D)-associated variants were enriched in CD4+ T cell caQTLs (naïve CD4+ T logOR=1.9, p=0.024; activated CD4+ T logOR=1.5, p=0.043), where T cells are the critical cell type in the pathogenesis of T1D^30^. Lymphocyte count-associated variants were enriched in caQTLs for lymphocyte cell types (memory B logOR=2.9, p=0.024; naïve CD4= T logOR=1.7, p=0.012). Strong enrichments for other traits may similarly point to cell types involved in trait biology. For example, child onset asthma-associated variants were enriched in activated CD4+ T cells (logOR=2.5, p=0.001) and memory B cells caQTLs (logOR=4.1, p=0.001) (**Figure 4c**), and ulcerative colitis-associated variants were strongly enriched in classical monocyte caQTLs (logOR=2.6, p=0.001).

Among fine-mapped variants that were immune cell caQTLs, 185 had a posterior probability >1% and were either in a distal site linked to a gene promoter or in a promoter site directly (**Figure 4d, Supplementary Table 9**). Among these, at multiple loci fine-mapped variant caQTLs replicated cell type-specific effects observed in previous studies^31,32^. For example, at the 5q11.2 locus associated with rheumatoid arthritis (RA), among the two candidate variants with highest causal probability rs28722705 (PPA=.70) and rs7731626 (PPA=0.28) only rs7731626 mapped in an accessible chromatin site (**Supplementary Figure 9a**). This variant was a caQTL in naive CD4+ T cells (π=0.38, q=0.06), and was co-accessible with the *IL6ST* and *ANKRD55* promoters (**Supplementary Figure 9b-e**). A previous study identified rs7731626 as likely causal for multiple sclerosis and RA and was also a T cell-specific eQTL for both *IL6ST* and *ANKRD55*^31^. At the 15q22.33 locus, rs17293632 was fine mapped in multiple traits including Crohn’s disease (PPA=.28) and asthma (PPA=.16), and was a monocyte caQTL (π=.40, q=.0138) which was co-accessible with the promoter of multiple SMAD3 isoforms (**Supplementary Figure 9f-i)**. The high effect C allele of rs17293632 was also predicted to have allele-specific binding to a JUN/FOS TF family motif, which were specifically enriched in monocyte caQTLs (**Supplementary Figure 9j, Figure 2g**).

In another example, at the 6q15 locus associated with type 1 diabetes (T1D) and multiple other traits, rs72928038 (PPA=.07) was a caQTL in naïve CD4+ T cells (π=0.26, q=4.9×10^−3^) where the reference and T1D-protective allele G had increased accessibility (**Figure 4e-f**). The site harboring rs72928038 was specific to naïve CD4+ T cells and was co-accessible with multiple gene promoters including *BACH2* (**Figure 4e, Supplementary Table 9**). The G allele was also predicted to have allele-specific binding to ETS family motifs, which were broadly enriched among T cell caQTLs (**Figure 4g**. We validated the allelic effects of this variant on regulatory activity using reporter assays in Jurkat T cells. There were significant effects on enhancer activity in luciferase gene reporter assays where the G allele had increased activity (Two-sided t-test, P=.015), and allele-specific transcription factor binding to the G allele in electrophoretic mobility shift assays (**Figure 4h**). Previous studies have shown that this variant is a QTL in CD4+ T cells^31^, and this site was linked to the *BACH2* promoter in promoter-capture Hi-C data in naïve CD4+ T cells^28^.

We next identified caQTLs for high-probability fine-mapped variants at loci without established molecular mechanisms. At the 11q23 locus associated with child-onset asthma, we fine-mapped a single variant rs12365699 to near-causality (PPA=.98) (**Figure 4i, Supplementary Table 9**). This variant was a caQTL in activated CD4+ T cells, effector CD8+ T cells and memory B cells, where the reference and risk-increasing allele G had higher accessibility (CD4+ T π=0.36, q=5.4×10^−4^, CD8+ T π=0.33, q=6.5×10^−3^, B π=0.35, q=0.073, respectively) and was linked to the promoter regions of multiple genes including *CXCR5* and *NLRX1*, the latter of which is ~300kb distal to the variant (**Figure 4i,j**). The G allele of rs12365699 was also predicted to have allele-specific binding for ZSCAN16 (**Figure 4k**). At the 12p13.33 locus associated with lymphocyte count, we also fine mapped a likely causal variant rs34038797 (PPA =.94), which had the same effect in all cell sub-types (strongest association in classical monocytes π=0.26, q=9.2×10^−4^) and was co-accessible with multiple genes (**Supplementary Figure 10a-c**). The C allele had higher accessibility and higher predicted affinity with ETS transcription factors, which were ubiquitously enriched in immune cell caQTLs (**Supplementary Figure 10d**).

## DISCUSSION

In this study we demonstrated that profiles derived from single nucleus ATAC-seq assays can be used to map chromatin accessibility QTLs in individual cell types, even with modest sample sizes. While we profiled only a small number of samples in our study, we identified thousands of immune cell type and sub-type caQTLs. A likely contributor to the large number of caQTLs we identified despite the small sample is the high depth at which samples were sequenced, which provides greater power for allelic imbalance mapping. Supporting this, we identified few caQTLs when performing population-based QTL mapping only. As the number of unique reads covering a variant can in theory be much higher for snATAC-seq compared to bulk ATAC-seq due to the thousands of unique libraries per assay, the value of snATAC-seq compared to bulk ATAC-seq in mapping allelic imbalance is even more pronounced. Deeply sequenced snATAC-seq assays even in few samples therefore represent an effective approach to map genetic effects on chromatin profiles from multiple cell types in a heterogeneous tissue.

Mapping caQTLs at cell type resolution enabled insights into cell type-specific regulation that are obscured from assays of bulk tissue chromatin. For example, we identified variants mapping in sites active in all cell types but with allelic effects on only a few cell types. We also identified examples of variants with opposite effects on different cell types resulting in no net effect in bulk. In both of these scenarios, simply annotating bulk caQTLs using reference maps of cell type-specific chromatin sites would not be sufficient to uncover these effects, and therefore requires mapping accessible chromatin profiles in each cell type directly. Single cell data also enabled additional cell type-specific analyses such as linking distal sites to putative target gene promoters using co-accessibility^27^. While high-resolution maps of distal 3D interactions exist for many immune cell types in promoter-capture Hi-C^28^, most other tissues do not currently have such cell type-resolved interaction maps and therefore cell type co-accessibility data will be particularly valuable in annotating distal caQTLs in these tissues.

Although we mapped thousands of immune cell type caQTLs from few snATAC-seq samples, our study design also has several notable limitations. Most importantly, there was a large difference in the number of caQTLs per cell type or sub-type dependent on the number of cells assayed. For example, we identified few significant caQTLs for the less common sub-types identified in our data such as adaptive NK cells and memory B cells. There are even further sub-divisions of immune cell types that we were not able to identify due to the resolution of snATAC-seq profiles. As identifying caQTLs from rarer cell types and sub-types will therefore require many additional snATAC-seq assays to sufficiently define their profiles, cell sorting may represent a more efficient and cost-effective strategy at present for QTL mapping in these cell types. An additional limitation of our study was that, due to the small number of samples profiled, we focused only on variants mapping in accessible chromatin sites directly in order to leverage allelic imbalance. As we did not test all variants at a locus for association to each site, we had limited ability to formally compare caQTL association and disease association signals, for example using colocalization techniques^33^. Moving forward studies profiling larger sample sizes and cell numbers will help circumvent these limitations. Furthermore, data from mutiomic assays of joint gene expression and accessible chromatin will help resolve cell types and sub-types and facilitate joint caQTL and eQTL mapping^34^.

In summary, we identified thousands of caQTLs in immune cell type and sub-types from peripheral blood samples using single cell chromatin accessibility assays. Immune cell caQTLs mapped to hundreds of loci associated with complex immune traits and disease and therefore represent a valuable resource for interpreting the molecular mechanisms of these loci. Given the ability to deconvolute individual cells into their sample-of-origin^20^, one promising strategy moving forward will be to pool samples prior to running snATAC-seq assays, which will reduce the per-sample cost and facilitate studies of greater genotype diversity. Mapping cell type-specific chromatin exposed to disease-relevant conditions and stimuli will also help uncover the breadth of genetic effects^23,35^. Together these efforts will enable more comprehensive annotation of variant function in human cell types and their contribution to complex disease.

## METHODS

### Single nuclei ATAC-Seq

Peripheral blood mononuclear cells (PBMCs) from 10 individuals (4 females and 6 males) were purchased from HemaCare (Northridge, CA) and profiled for snATAC using 10x Genomics Chromium Single Cell ATAC Solution, following manufacturer’s instructions (Chromium SingleCell ATAC ReagentKits UserGuide CG000209, Rev A) as described previously^17^. Briefly, cryopreserved PBMC samples were thawed, resuspended in 1 mL PBS (with 0.04% FBS) and filtered with 50 μm CellTrics. Cells were centrifuged and permeabilized with 100 μl of chilled lysis buffer (10 mM Tris-HCl pH 7.4, 10 mM NaCl, 3 mM MgCl2, 0.1% Tween-20, 0.1% IGEPAL-CA630, 0.01% digitonin and 1% BSA) for 3 min on ice and then washed with 1mL chilled wash buffer (10 mM Tris-HCl pH 7.4, 10 mM NaCl, 3 mM MgCl2, 0.1% Tween-20 and 1% BSA). After centrifugation, pellets were resuspended in 100 μL of chilled Nuclei buffer (2000153, 10x Genomics) in a final concentration of 3,000 to 7,000 of nuclei per μl. 15,300 nuclei (targeting 10,000) were used for each sample. Tagmentation was performed using nuclei diluted to 5 μl with 1X Nuclei buffer, 10x ATAC buffer and ATAC enzyme from 10x Genomics, for 60 min at 37°C. Single cell ATAC-seq libraries were generated using the Chromium Chip E Single Cell ATAC kit (10x Genomics, 1000086) and indexes (Chromium i7 Multiplex Kit N, Set A, 10x Genomics, 1000084) following manufacturer instructions. Samples were sequenced to an average depth of 178 million 50-nt read pairs each, using an illumina HiSeq4000 instrument at the UCSD Institute for Genomic Medicine. Alignment to the hg19 genome and initial processing were performed using the 10x Genomics Cell Ranger ATAC v1.1 pipeline. We filtered reads with MAPQ<30, secondary or unmapped reads, and duplicate reads from the resulting bam files using samtools^36^. Sample information and a summary of the Cell Ranger ATAC-seq quality metrics are provided in **Supplementary Table 1**.

### Quality control, clustering and cell type assignment

For each sample we performed multiplet removal (N_cells_=1,311) using Cell Ranger’s custom multiplet removal script (version 1.1). The genome was split into 5 kb windows and windows overlapping blacklisted regions from ENCODE (version 2) were removed. For each sample, a sparse *m* x *n* matrix containing read depth for *m* cells (identified using the snATAC-seq barcodes) passing read depth thresholds at *n* windows was then generated.

Initial cell clustering was performed separately for each snATAC-seq sample sparse matrix using scanpy (version 1.5). Highly variable windows were extracted using mean read depths and dispersion was normalized. Read depth was normalized, and the log-transformed read depth was regressed out for each cell. Principal component analysis was then performed, and the top 50 principal components were used to calculate the nearest 30 neighbors using the cosine metric. This cosine metric was then used to perform UMAP dimensionality reduction clustering with the parameters ‘min_dist=0.3’, along with further sub-clustering using the Louvain clustering algorithm with the parameters ‘resolution=1.25’. Clusters with a low fraction of reads in promoter, a low log usable read count, and/or a low fraction of reads in peaks were iteratively removed for each sample (N_cells_=6,333). The samples filtered for low quality cells were then merged, and PCs and UMAP dimensions were obtained as above and Harmony was used to correct for donor batch effects^37^. Manual doublet removal was then performed by removing Louvain-defined sub-clusters that had higher than average useable read counts, mapped between clusters and/or expressed multiple marker genes. Clusters that did not have uniform representation across samples were also removed. A total of 14,268 cells were removed during all of the quality control steps. UMAP dimensionality reduction was performed again using the same parameters on the remaining cells in order to re-cluster a final time.

In order to assign cell type and sub-type identity to each cluster, we determined chromatin accessibility at 5 kb windows around promoter regions of known marker genes (see **Supplementary Table 3**).

### Peak Calling

For each cluster mapped reads were extracted from all cells within the cluster. Reads aligning to the positive strand were shifted by +4 bp and reads aligned to the negative strand were shifted by −5 bp. Reads were extended to 200 bp and then centered, and bed files were created from the resulting read coordinates. We then called peaks with MACS2^22^ from the bed files using the parameters ‘-q 0.05’, ‘-nomodel’, ‘-keep-dup all’, ‘-g hs’, and ‘-B’. The read count pileup bedgraph was sorted and normalized to counts per million (CPM), converted to bigwig and visualized using the UCSC Genome Browser. We created a merged peak set by combining narrow peak files across all cell type and sub-type clusters into a single bed file.

### Comparison with bulk immune cell ATAC-seq data

We obtained published data of FACS-sorted immune cell types (GSE118189)^23^, mapped reads to hg19 using bwa mem^38^ and removed duplicate reads. We merged replicate samples and performed peak calling for each cell type as described above. Mapped reads from immune cell types and sub-types derived from snATAC-seq in this study and from the bulk immune cell ATAC-seq profiles were used to generate bedgraph files using bedtools^39^. Read counts were normalized to CPM and bigwig files were generated using ENCODE ‘bedgraphToBigWig’^40^. We created a bed file of the union of peak calls from snATAC-seq and bulk ATAC-seq using bedtools. We then compared bulk ATAC-seq cell type and snATAC-seq cell type normalized read count profiles within the union peak set using deeptools ‘multiBigWigSummary’^41^. A heatmap of the clusters of Spearman rank correlation coefficients indicating similarity between bigwig files was generated using deeptools ‘plotCorrelation’ and the summary comparison from ‘multiBigWigSummary’.

### Sample genotyping and imputation

Genomic DNA form PBMC samples was extracted using the PureLink genomic DNA kit (Invitrogen). Genotyping was performed using Infinium Omni2.5-8 arrays (Illumina) at the UCSD Institute for Genomic Medicine. Genotypes were assigned with GenomeStudio (v.2.0.4) with default settings. Variants with minor allele frequency (MAF) < 0.01 or with ambiguous alleles (G/C, or A/T) and MAF > 0.4 were filtered out using PLINK. For the remaining variants, we imputed genotypes into the Haplotype Reference Consortium (HRC) r1.1 panel using the Michigan Imputation Server with minimac4. We then retained variants with imputation quality R2>0.7

### Identification of chromatin accessibility QTLs

For each sample, we split reads in the snATAC .bam files according to cluster label. For each cell type and sub-type cluster, we generated peak count matrices (peak x sample) using merged peak site coordinates and the split .bam files using featureCounts^42^. We then obtained VCF files of SNPs located within peaks and annotated allelic read counts using RASQUAL tools^24^. We filtered for variants heterozygous in at least 2 samples. For the ‘bulk’ experiment we ignored cell type labels and used all reads.

For each cell type and sub-type, we retained only accessible sites with at least 5 reads on average across samples. To perform caQTL analysis we used RASQUAL and tested for association between each peak and variants contained in the peak itself or in other peaks within a 10Kb window. We included the library size of each sample calculated using the rasqualCalculateSampleOffsets() function and read count covariates using make_covariates() function in each model. The number of ATAC-seq read count covariates were dynamically calculated for each cell type and sub-type and therefore different cell types/sub-types had different numbers of covariates. We also included the first four principal components derived from genotype data together with major 1KG populations as covariates in each model.

For each peak, we calculated adjusted p-values accounting for the number of variants tested per peak, and the variant with the minimum adjusted p-value was marked as the lead variant. To correct for multiple testing genome-wide, we performed permutations of labels across samples and counts across alleles of heterozygous variants. For the permutations across samples, we required that the labels were swapped within the samples of European and American ancestry separately. We then repeated the association tests and calculated an empirical FDR (10%) by comparing the q-values of the real and permuted association results.

To estimate the correlation of effect sizes of caQTLs across cell types and ‘bulk’ data we calculated the spearman correlation coefficient of effect sizes (π) in each pair of cell types and “bulk”. For each comparison we selected lead SNP-peak pairs that were significant caQTLs in at least one of the two cell types. Correlation coefficients were tabulated in a matrix and hierarchically clustered using ‘pheatmap’. Bulk-like caQTLs were compared with caQTLs from 24 LCLs also calculated using RASQUAL^24^. Of 172,241 peaks tested in PBMCs, 65,787 intersected with a peak tested in LCLs, and 660 were caQTLs in both dataset (FDR 10%). Enrichment was estimated using Fisher’s exact test. To calculate coordination of caQTLs effects we restricted the analysis to those peaks having the same lead variants (589), 164 of which were caQTLs in both datasets. Monocyte and CD4+ and CD8+ T-cells single-cell caQTLs were compared with H3K27ac QTLs and eQTLs from FACS sorted Monocytes and T-cells from the BLUEPRINT project, calculated using WASP and the Combined Haplotype Test at FDR 10%, which similarly to RASQUAL takes into account both allelic and population effects. For each comparison we selected variants tested in both datasets and calculated enrichment for shared variant QTLs (lead variants only) using Fisher’s exact test.

### Transcription factor motif analysis

To identify enriched motifs that were altered by caQTLs we used the package MotifBreakR^25^. First, we selected 109,554 SNPs that were tested in any of the cell types for caQTLs and imported them using the function snps.from.file(), using hg19 as reference genome. Then we determined if they disrupted TF motifs from the HOCOMOCO v10 human database^26^, comprising 640 motifs corresponding to 595 unique TFs, and accessed via MotifDb. The following motifbreakR() function parameters were used: filterp = TRUE, method=“ic”, = 5e-4, BPPARAM = BiocParallel::bpparam(“SerialParam”). SNPs that resulted in disruption of any TF motif with a *strong* effect (defined by motifbreakR) were considered as motif altering (n=107,280). To calculate enrichment for alteration of specific TF in caQTLs of individual cell types (B-cell, CD4_T-cell, CD8_T-cell, NK_cell, Monocyte), we performed a one-tailed exact binomial test (binom.test(alternative= “greater)) comparing the frequency of alteration of a motif by caQTLs to the total frequency of motif alteration in the tested SNPs for each cell type. Significant enrichment was considered at a Benjamini & Hochberg corrected P-value<0.05. To display enriched motifs we used the packages MotIV and motifPiles, selecting the top motifs in each cell type ranked by p-value.

### Single cell co-accessibility

Peak-to-peak co-accessibility was calculated using Cicero (version 1.1.5)^27^ for B-cells, CD4+ T-cells, CD8+ T-cells, NK cells, and Monocytes. We created a sparse binary matrix encoding the snATAC-seq barcodes for each cell in a given cell type and the superset of ATAC-seq peaks for all cell types, indicating which cells were accessible in which peaks. For each cell type, the cicero function ‘make_cicero_cds’ was used to aggregate cells into bins of 30 nearest neighbors (parameter k=30) from the UMAP reduced dimensions obtained from clustering. We then calculated co-accessibility scores using a window size of 1 Mb. Once co-accessibility scores were calculated, a threshold of 0.05 and a minimum distance of 10 kb were used to define pairs co-accessible for a given cell type. We also generated cell type co-accessibility for each sample individually and only retained sites co-accessible at .05 in at least two individual samples. A peak was categorized as ‘promoter’ if it fell within a 2 kb window of a transcription start site based on GENCODE (version 19) promoter annotations^43^, and otherwise was categorized as ‘distal’.

To validate the cell-type specificity of promoter-distal sites connections calculated using co-accessibility, we compared them to chromatin interactions from promoter capture Hi-C (pCHi-C) data previously generated in 16 immune cell types and sub-types^28^. We obtained the list of promoter baits and the matrix containing CHiCAGO scores for all interactions in all immune cell type. First, for each pair of peaks that we analyzed in each cell type, we filter those where at least one peak intersected (+/- 1kb) a pCHi-C bait using pgltools^44^. Then, we identified overlapping connection between the filtered pairs of sites in each of our 5 cell type (B-cells, CD4+ T-cells, CD8+ T-cells, NK cells, and Monocytes) and the pCHi-C connection (CHiCAGO score >= 5) from each of the 16 blueprint adult cell types (Mon, Mac0, Mac1, ‘Mac2’, Neu, MK, EP, Ery, nCD4, tCD4, aCD4, naCD4, nCD8, tCD8, nB, tB) using the function compare_connections() from the Cicero package. For each celltype-celltype comparison we then estimated the enrichment for the co-accessible sites in pCHi-C connection using Fisher’s exact test. Odds ratios for each comparison were tabulated and displayed using heatmap. For each of the matching cell types (B-cells-tB, CD4+T-cells-tCD4, CD8+T-cells-’tCD8’, and Monocytes-Mon) we also calculated enrichment at different peak distances (10-50, 50-100, 100-200, 200-350, 350-1000 kb).

### Distal effect of caQTL variants on coaccessible-promoters

To examine the effect of caQTLs on co-accessible sites, for each of the 5 major cell types we took the lead caQTL variant and tested for association with accessibility level of the co-accessible site using RASQUAL^24^. We used the same method as above for caQTLs with the exception of adopting a more relaxed FDR threshold of 20% instead of 10%. caQTLs-coaccessible peaks were then filtered to retain only enhancer-promoters and promoter-promoter co-accessible peaks (B-cells n=828, CD4+ T-cells n=2,323, CD8+ T-cells n=1,909, NK cells n=1,489, Monocytes n=2,243, with an average number of 3.45 co-accessible promoters for each caQTL). Pearson correlation of effect sizes was calculated between variant effect on the original caQTL peak and on one of the co-accessible promoters (with lowest RASQUAL p-value of association), and only considering co-accessible peaks at >10kb of distance.

### Genetic fine mapping analysis

We obtained genome-wide summary statistics for immune-related phenotypes including blood cell type counts^45^, autoimmune diseases^46–50^, and inflammatory diseases^51–53^. For each study, we obtained lists of index variants for each independent signal from the supplement. We used PLINK^54^ to estimate linkage disequilibrium (LD) between these index variants and all variants within ±2.5 Mb using samples of European ancestry from the 1000 Genomes Project^55^. For each signal, we first pre-filtered variants in at least low LD (r^2^>0.1) with the index variants. We calculated approximate Bayes factors^56^ (aBF) for each variant using the effect estimates (β) and standard errors (SE), assuming prior variance w=0.04. We calculated the posterior probability of association (PPA) by dividing the aBF for each variant by the sum of aBFs for all variants included in the signal. We then defined the 99% credible set as the smallest set of variants that added up to 99% PPA. Fine-mapped variants were annotated using cell type and sub-type caQTLs, considering each lead variant as well as variants with the same q-value of the lead variant for each caQTL. Fine-mapped caQTL variants with PPA>1% were then further annotated with co-accessible promoters (**Supplementary Table 8**).

To test for enrichment of caQTLs for complex immune traits we calculated the cumulative PPA of variants overlapping immune cell sub-type caQTL peaks across all credible sets for each trait. For each cell sub-type, we defined a background set of peaks tested for association but did not have significant caQTLs. We estimated an empirical distribution for the total PPA using 1,000 random draws of peaks from the background equal in number to the caQTL sites. For each test (trait vs cell sub-type) a p-value was calculated by comparing the total PPA within caQTL peaks to the empirical distribution.

### Luciferase gene reporter assays

Human DNA sequences (Coriell) with reference allele for rs72928038 (*BACH2* intron) were cloned in forward orientation in the luciferase reporter vector pGL4.23 (Promega) using the primers: forward, AGCTAGGTACCACACTCAGTGGTTGGGGTTT, and reverse, TACCAGAGCTCCTGGATAGAGGTCCCAGTCG and the enzymes SacI and KpnI. Alternate allele plasmids were generated via site directed mutagenesis (Q5 SDM kit, New England Biolabs) using the following primers: forward, CGGATTTCCTaTAAGCTGATC, reverse, TCCCTATTTGTGTGTAATG.

Jurkat cells were maintained in culture at a concentration of 1×10^05^/mL-1×10^06^/mL. Approximately 0.5×10^06^ cells per replicate (3 replicates) were co-transfected with 500 ng of firefly luciferase vector containing either the reference or alternate allele or an empty pGL4.23 vector as a control, and 50 ng pRL-SV40 Renilla luciferase vector (Promega), using the Lipofectamine LTX reagent. Cells were collected 48 hours post transfection and assayed using the Dual-Luciferase Reporter system (Promega). Firefly activity was normalized to the Renilla activity and expressed as fold change compared to the luciferase activity of the empty vector (RLU). A two-sided t-test was used to compare the luciferase activity between the two alleles.

### Electrophoretic mobility shift assays

EMSAs were performed according to manufacturer’s instruction, with changes indicated below, using the LightShift™ Chemiluminescent EMSA Kit (Thermo Scientific, 20148). Biotinylated and non-biotinylated single-stranded oligonucleotides harboring the rs72928038 variant (5’-TAGGGACGGATTTCCTGTAAGCTGATCTTGAAG-3’, 5’-TAGGGACGGATTTCCTATAAGCTGATCTTGAAG-3’) were purchased from Integrated DNA Technologies. Nuclear extract from E6-1 Jurkat T cells (ATCC TIB-152), cultured as described above, was obtained using the NE-PER Nuclear and Cytoplasmic Extraction Reagents (Thermo Scientific, 78833). Binding reactions were carried in a total volume of 20 μl, with 10X Binding Buffer (100 mM Tris pH 7.5, 500 mM KCl and 10 mM DTT), 2.5% glycerol, 5 mM MgCl2, 0.05% NP40, 50 ng Poly(dI:dC), 100 fmole of biotin-labeled probe, and 5.1 μg nuclear extract. For competition experiments, 20 pmol of unlabeled probe was added. Competition reactions were incubated at room temperature for 10 mins before the addition of the biotin-labeled probe. At the addition of the biotin-labeled probe, all reactions were incubated at room temperature for 20 min. Reactions were loaded onto a 6% polyacrylamide 0.5X TBE Gel (Invitrogen, EC62655BOX) for electrophoresis and transferred for 45 mins to a Biodyne™ B Pre-Cut Modified Nylon Membrane, 0.45μm (Thermo Scientific, 77016). Transferred DNA was UV-crosslinked for 15 mins, and the biotinylated probes were detected using Chemiluminescent Nucleic Acid Detection Module (Thermo Scientific, 89880) following the manufacturers instruction, with initial blocking increased to 60 mins. The image was acquired using C-DiGit Blot scanner (LI-COR Biosciences, Model 3600).

## SUPPLEMENTARY FIGURE LEGENDS

**Supplementary Figure 1. Population structure of PBMC samples.** The first four principal components derived from joint analysis of genotype data from the 1000 Genomes Project and PBMC samples. Samples in 1000 Genomes are colored by major population group, and the PBMC samples are colored in pink.

**Supplementary Figure 2. Defining immune cell types and sub-types from snATAC-seq profiles.** a) UMAP plots showing promoter accessibility in a 1 kb window around the TSS for selected cell type marker genes (see Supplementary Table 3). b) Genome browser plots showing aggregate read density (scaled to uniform 1×10^5^ read depth, range: 5-35, shown on vertical axis for each plot) for cells within each cell type for selected cell type marker genes.

**Supplementary Figure 3. Immune cell type snATAC-seq profiles in individual PBMC samples.** a) UMAP plot showing cells clustering in each of the 10 PBMC samples assayed in this study. b) Scatter plot comparing cell type proportions obtained from cluster analysis versus those obtained from flow cytometry, excluding leukocytes. Proportions represent the fraction of all cells in each sample (see Supplementary Table 4 for individual sample proportions). Each dot represents an individual sample. c) Barplot showing the number of cells assigned to 14 distinct immune cell types and sub-types in each sample. d) Barplot showing the relative proportion of cells from each sample in each immune cell type and sub-type.

**Supplementary Figure 4. Comparison of ATAC-seq peaks from PBMC snATAC and FACS sorted PBMCs.** Heatmaps and hierarchical clustering of Spearman correlation coefficients for pairwise comparisons of genome-wide ATAC-seq profiles across (a) cell-types or (b) sub-types from PBMC snATAC from this study (in blue) and from a published bulk ATAC-Seq study using FACS sorted immune cells (in black).

**Supplementary Figure 5. Comparison of caQTL effects with and without the allelic imbalance component.** For each cell type (first 5 plots) and cell sub-type (remaining 10 plots), a scatter plot show the consistency between caQTL effect considering both allelic and population effect (x-axis) and the effect for the same variant-peak pair using only the population component (y-axis) obtained running RASQUAL using the --population-only option. The percentage of discordant effect are indicated.

**Supplementary Figure 6. Comparison of snATAC-seq caQTLs with LCL caQTLs from 24 individuals.** a-e) Significant PBMC caQTLs in a) B-cells, b) CD4+ T-cells, c) CD8+ T-cells, d) Monocytes and e) NK-cells and their overlap with caQTLs from LCLs (Venn diagram, considering only peaks tested in both datasets. For each shared caQTLs between each cell type and LCLs, a scatter plot shows effect sizes for caQTLs found in both studies and having the same lead variant. f) Table with p-values and odds ratio from two-tailed Fisher’s exact test for enrichment of cell-type caQTLs in LCLs caQTLs.

**Supplementary Figure 7. Examples of caQTLs specific to immune sub-types.** a) Examples of cell-type specific caQTLs due to presence of the peak in a single cell type. From left to right, rs1957554 is a caQTL for a naïve B cell-specific site, rs7094953 is a caQTL for a naïve CD4+ T cell-specific site, rs3014874 is a caQTL for a classical monocyte-specific site, and rs59176853 is a caQTL for a cytotoxic NK cell-specific site. Top panels: colored-coded box-plots show association in the different cell types, white box-plots show corresponding caQTL in bulk PBMCs. Association q-values are shown on the top and variant genomic location (hg19) is shown at the bottom. Bottom panels: genome-browser screenshot of snATAC-seq in different cell types. b) Example of caQTL specific to classical monocytes (rs747748) although the peak is active in all immune cell types. Box-plots show association with the same variant in 10 immune sub-types and the bulk PBMCs. Right: genome-browser screenshot of cell-type color-coded snATAC-seq peaks and position of the variant. c) Example of a caQTL specific to effector CD8+ T cells (rs61943586) although the peak is active in all cell types.

**Supplementary Figure 8. Correlation of caQTL effects with effects at distal promoters across different distances**. Correlation in the effects of caQTL variants on the QTL site and co-accessible promoter sites in each cell type, grouped by distance between the QTL site and co-accessible promoter site. Pearson correlation coefficient and number of co-accessible pairs of peaks are indicated.

**Supplementary Figure 9. Additional examples of immune cell type caQTLs at fine-mapped complex immune trait loci with cell-specific effect.** a) Regional plot of a locus on chr5 associated with rheumatoid arthritis, with the four credible set variants highlighted in red. The candidate causal variant rs7731626 is indicated with a triangle and PPA. b) Chromatin signal in naïve CD4+ T cells in the region and co-accessible link between the site harboring rs7731626 and the *IL6ST* (aka *GP130*) promoter. The other three peaks co-accessible with rs7731626 map to the promoter of a non-coding isoform of *ANKRD55* (in green). c) Zoomed-in chromatin signals at the rs7731626 and the *IL6ST* promoter sites in all cell subtypes, showing specificity for naïve CD4+ T cells. d) Chromatin signal at the rs7731626 variant grouped by rs7731626 genotype in bulk PBMCs and naïve CD4+ cells and q-values of association. e) Top four predicted TF sequence motifs rs7731626, where the variant base is highlighted. Twelve other motifs predicted to be altered are not shown. f) Regional plot of the locus on chr15 associated with Crohn’s disease, with the five credible set variants highlighted in red. The candidate causal variant rs17293632 is indicated with a triangle and PPA. g) Chromatin signal in classical monocytes in the region and co-accessible link between the intronic enhancer harboring rs17293632 and three alternative *SMAD3* promoters. h) Zoomed-in chromatin signals at the rs17293632 (right) and *SMAD3* promoter sites in all cell subtypes, showing specificity for monocytes at the enhancer site and the two closest promoters. i) Chromatin signal at the enhancer site grouped by rs17293632 genotype in bulk PBMCs, monocytes (including all subtypes), classical monocytes and non-classical monocytes, with q-values of association (RASQUAL). j) Four predicted TF sequence motifs rs7731626, where the variant base is highlighted. Ten other similar motifs (ETS family) predicted to be altered are not shown.

**Supplementary Figure 10. Additional examples of immune cell type caQTLs at fine-mapped complex immune trait loci with high causal probability.** a) Regional plot of the locus on chr12 in the *NINJ2* gene showing association with Lymphocyte count, with the eight credible set variants highlighted in red. The candidate causal variant rs34038797 is indicated with a triangle and its PPA is shown. b) Chromatin signal in memory CD8+ T cells and Classical Monocytes in the same region and the co-accessible link between the site harboring rs34038797 and promoters of *CCDC77, WNK1, RAD52* (CD8+ T), *NINJ2* and *SLC6A12* (Monocyte). c) Chromatin signal at the rs34038797 variant grouped by rs34038797 genotype in all cell sub-types and q-values of association (RASQUAL) and zoomed-in genome browser track. d) Top five predicted TF sequence motifs altered by rs34038797, where the variant base is highlighted.

## SUPPLEMENTARY TABLES

**Supplementary Table 1. Summary of PBMC samples information.** For each of the 10 samples analyzed, sample name, lot number, donor ID, donor age, gender, ethnicity, blood type, and flow cytometry markers percentages are indicated.

**Supplementary Table 2. Summary of snATAC-seq cell ranger statistics.** For each of the 10 samples analyzed, we indicate snATAC-seq sequencing and mapping statistics.

**Supplementary Table 3. Marker genes references.** List of marker genes used to assign clusters to PBMC cell types and sub-types and corresponding reference papers.

**Supplementary Table 4. Clustering vs. flow cytometry cell type proportions.** Comparison between cell type proportions in each sample as estimated by flow cytometry and snATAC.

**Supplementary Table 5. Immune cell type and sub-type accessible chromatin sites.** Merged bed file of all ATAC peaks sites called in each cell type and sub-type, used for all analyses.

**Supplementary Table 6. Immune cell type and sub-type caQTLs.** RASQUAL results for all caQTLs significant at FDR 10% in each cell type (5 cell types and 10 sub-types) and pseudo-bulk analyses. The first 25 columns are outputs from RASQUAL, and p-values and q-values were calculated from columns 11 and 10, respectively.

**Supplementary Table 7. Transcription factor motifs enriched in immune cell type caQTLs.** List of TF motifs from the HOCOMOCO v.10 human database that were tested for enrichment in caQTLs and results of binomial test.

**Supplementary Table 8. Complex immune traits and diseases included in fine-mapping.** List of traits and corresponding GWAS study used for fine mapping.

**Supplementary Table 9. Immune cell type and sub-type QTLs at fine-mapped variants.** List of SNPs in credible sets for blood and auto-immune traits that are caQTLs in one or more cell types, have PPA >0.01 and are located either in gene promoters or in enhancers that are co-accessible with distal promoters. For each fine-mapped variant we report caQTL results and co-accessible promoters (multiple entries) in each of the cell types with significant caQTLs.

## SUPPLEMENTARY DATA FILES

**Supplementary Data 1. Summary statistics of caQTLs in PBMC cell types, subtypes, and bulk like data.** RASQUAL results for all peaks tested in each cell type (5 cell types and 10 sub-types) and pseudo-bulk analyses. The first 25 columns are outputs from RASQUAL, and p-values and q-values were calculated from columns 11 and 10, respectively.

**Supplementary Data 2. Fine-mapping credible sets for loci associated with 16 complex immune traits.** The 99% credible sets derived from fine-mapping of loci associated with 16 complex immune traits and disease.

## Notes

### Competing Interest Statement

KJG does consulting for Genentech and holds stock in Vertex Pharmaceuticals

