## Supplemental Figures for "Mapping genetic effects on cell type-specific chromatin accessibility and annotating complex trait variants using single nucleus ATAC-seq"

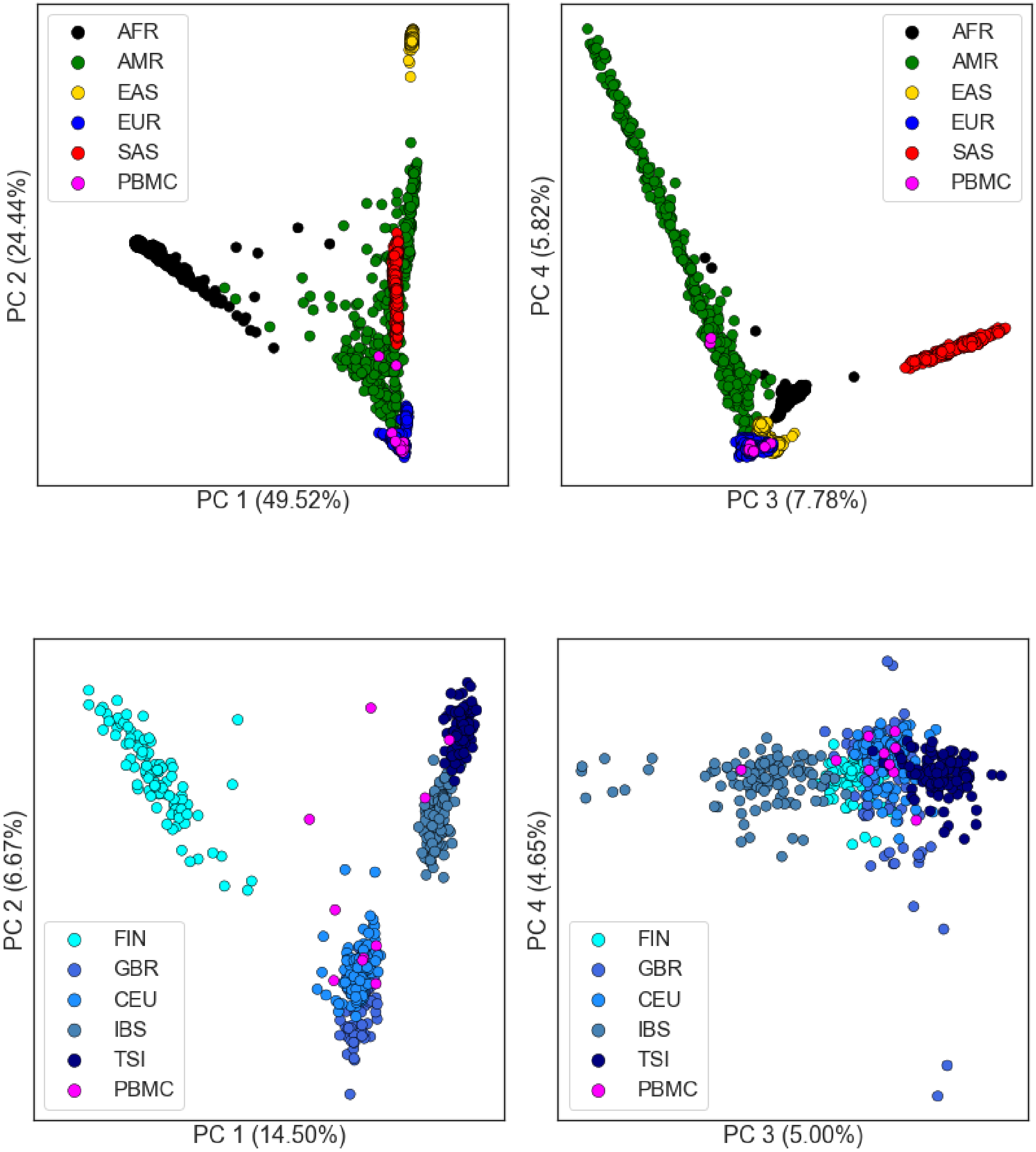
AFR
AMR
EAS
EUR
SAS
PBMC
PC 2 (24.44%)
PC 1 (49.52%)
PC 4 (5.82%)
PC 3 (7.78%)
FIN
GBR
CEU
IBS
TSI
PBMC
PC 2 (6.67%)
PC 1 (14.50%)
PC 4 (4.65%)
PC 3 (5.00%)

A

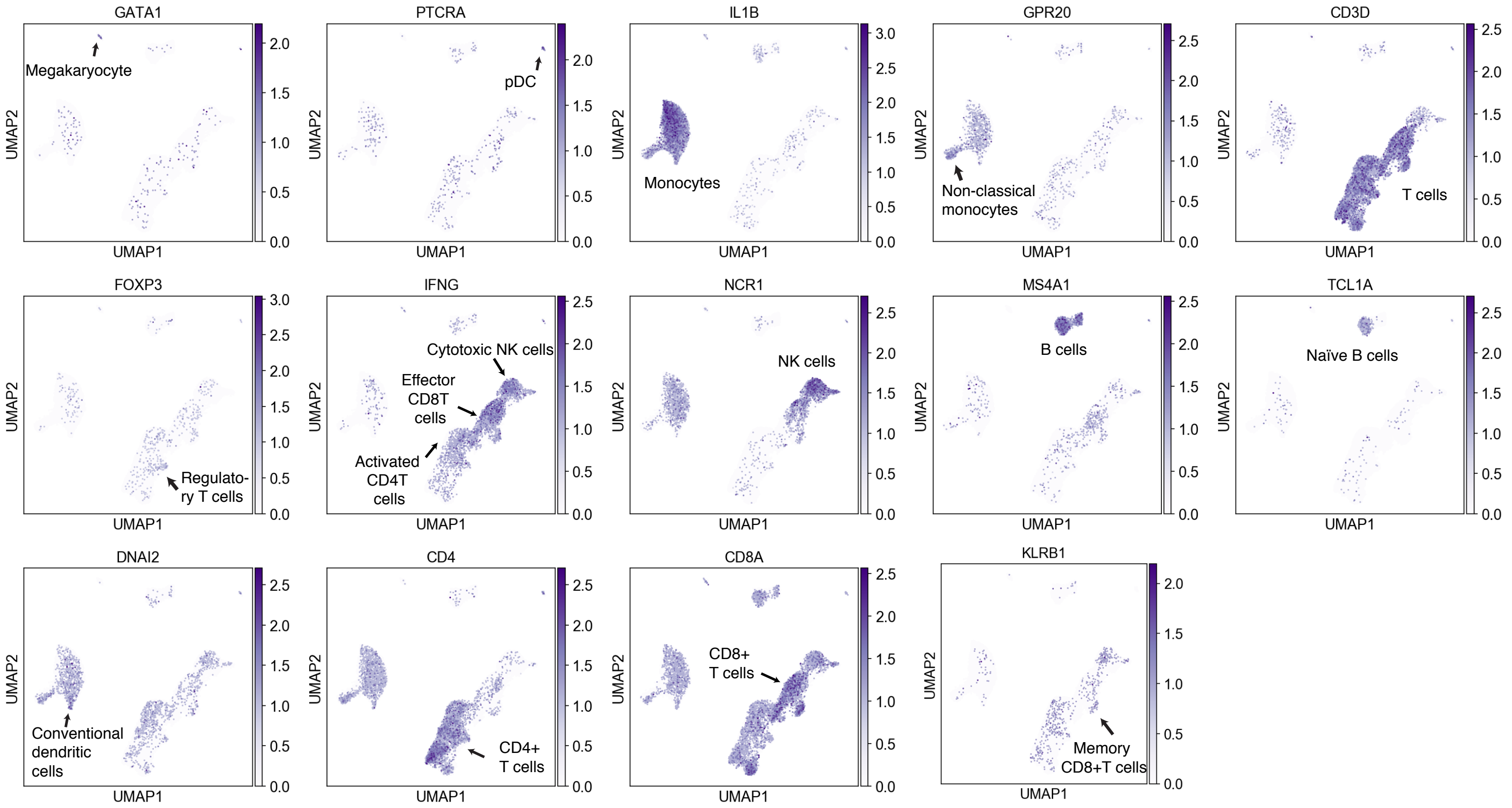
GATA1
PTCRA
IL1B
GPR20
CD3D
FOXP3
IFNG
NCR1
MS4A1
TCL1A
DNAI2
CD4
CD8A
KLRB1
UMAP1
UMAP2
Megakaryocyte
pDC
Monocytes
Non-classical monocytes
T cells
Regulatory T cells
Cytotoxic NK cells
Effector CD8T cells
Activated CD4T cells
NK cells
B cells
Naïve B cells
Conventional dendritic cells
CD4+ T cells
CD8+ T cells
Memory CD8+T cells

B

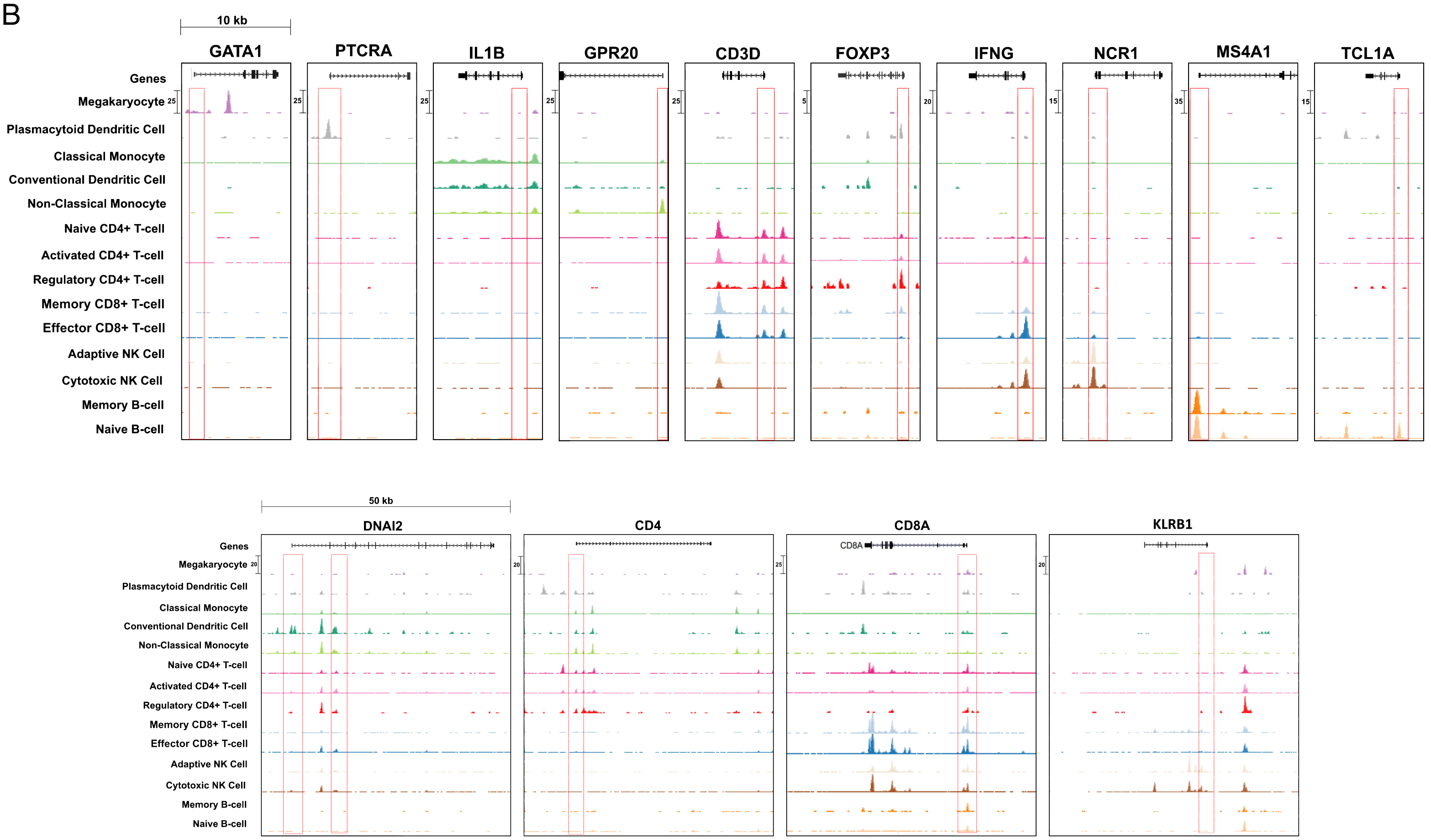
10 kb
GATA1
PTCRA
IL1B
GPR20
CD3D
FOXP3
IFNG
NCR1
MS4A1
TCL1A
Genes
Megakaryocyte
Plasmacytoid Dendritic Cell
Classical Monocyte
Conventional Dendritic Cell
Non-Classical Monocyte
Naive CD4+ T-cell
Activated CD4+ T-cell
Regulatory CD4+ T-cell
Memory CD8+ T-cell
Effector CD8+ T-cell
Adaptive NK Cell
Cytotoxic NK Cell
Memory B-cell
Naive B-cell
50 kb
DNAI2
CD4
CD8A
KLRB1

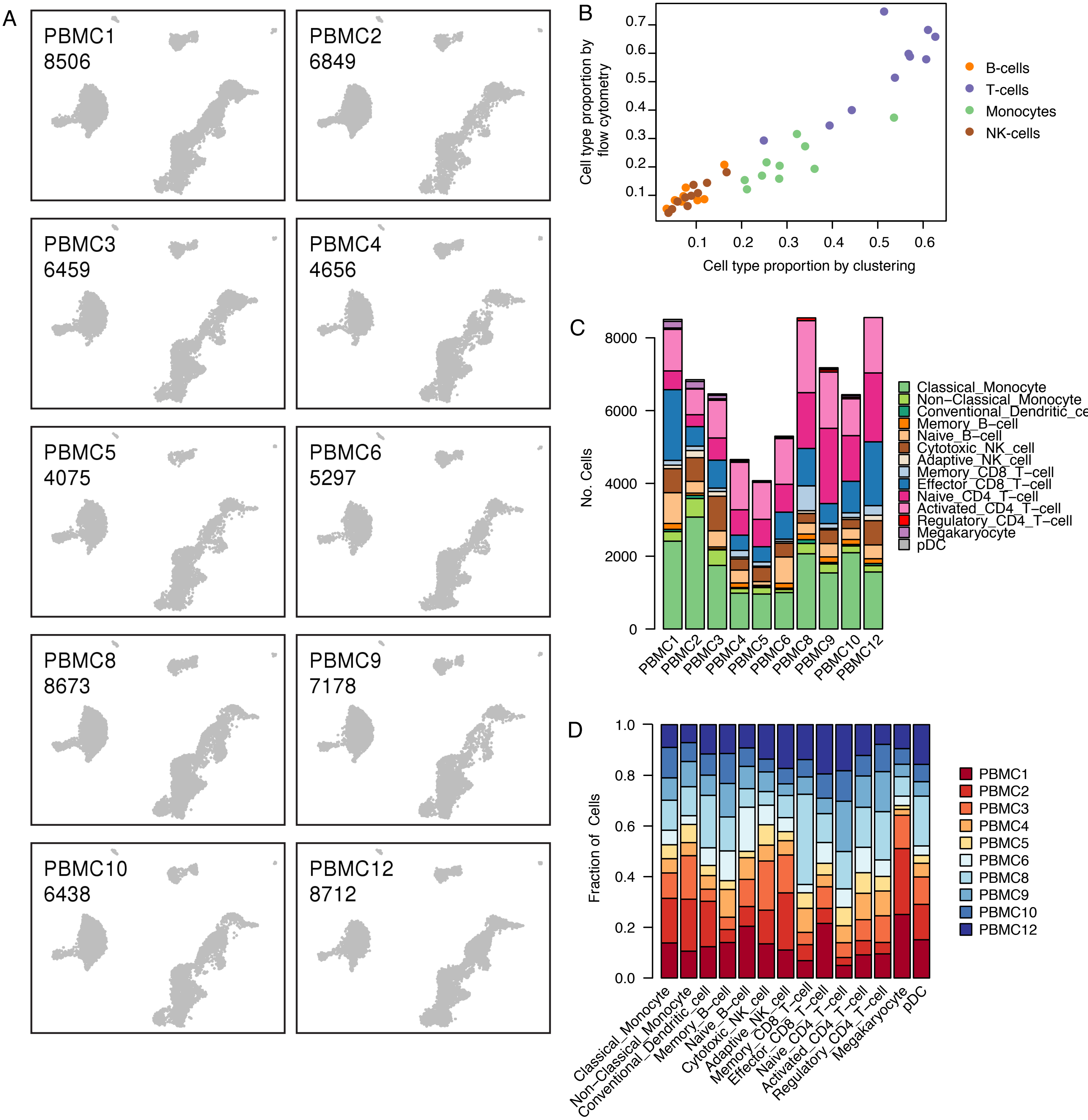
A
PBMC1
8506
PBMC2
6849
PBMC3
6459
PBMC4
4656
PBMC5
4075
PBMC6
5297
PBMC8
8673
PBMC9
7178
PBMC10
6438
PBMC12
8712
B
Cell type proportion by flow cytometry
Cell type proportion by clustering
B-cells
T-cells
Monocytes
NK-cells
C
No. Cells
Classical_Monocyte
Non−Classical_Monocyte
Memory_B−cell
Naive_B−cell
Cytotoxic_NK_cell
Adaptive_NK_cell
Memory_CD8_T−cell
Effector_CD8_T−cell
Naive_CD4_T−cell
Activated_CD4_T−cell
Regulatory_CD4_T−cell
Megakaryocyte
pDC
D
Fraction of Cells
Classical_Monocyte
Non−Classical_Monocyte
Conventional_Dendritic_cell
Memory_B−cell
Naive_B−cell
Cytotoxic_NK_cell
Adaptive_NK_cell
Memory_CD8_T−cell
Effector_CD8_T−cell
Naive_CD4_T−cell
Activated_CD4_T−cell
Regulatory_CD4_T−cell
Megakaryocyte
pDC
PBMC1
PBMC2
PBMC3
PBMC4
PBMC5
PBMC6
PBMC8
PBMC9
PBMC10
PBMC12

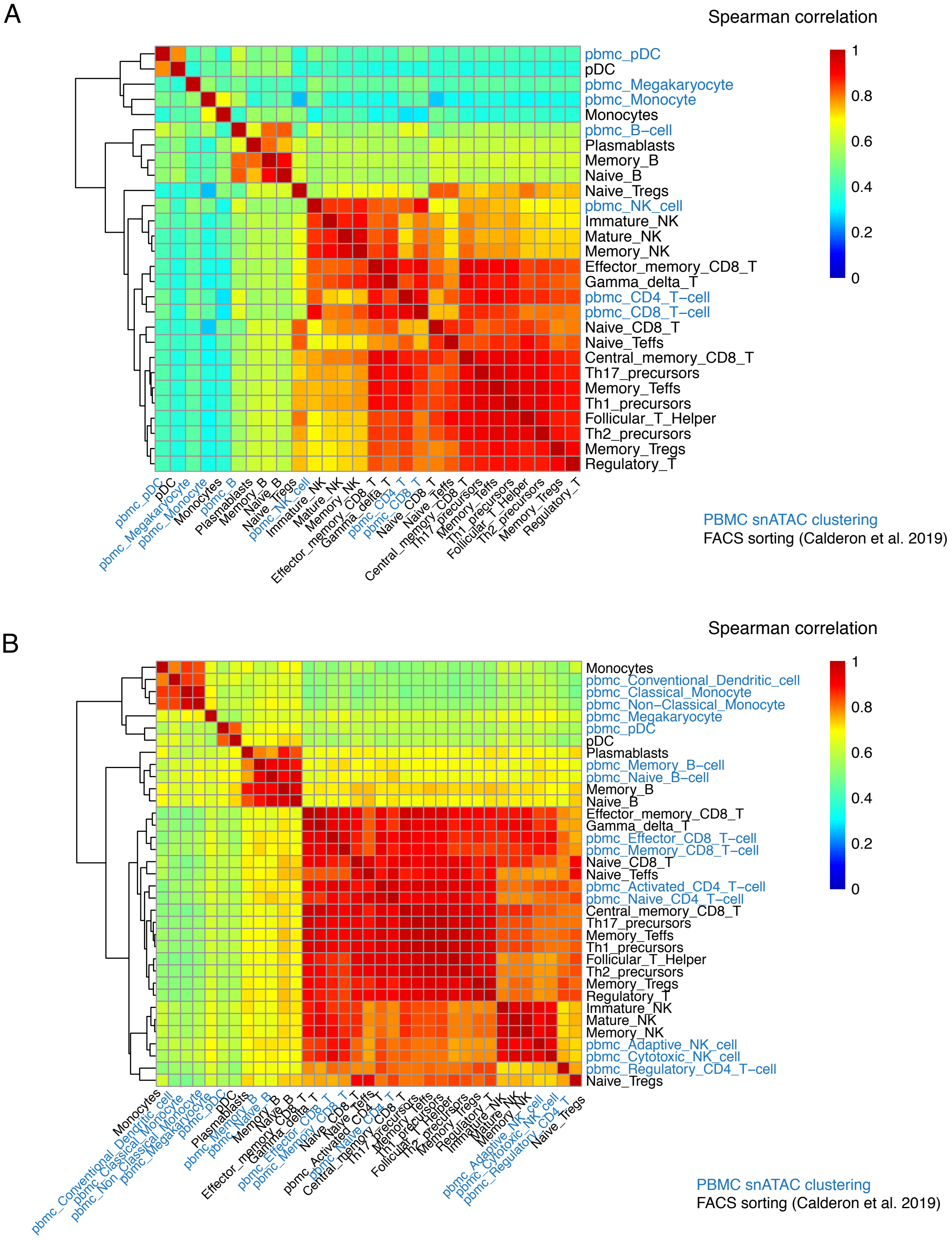
A
Spearman correlation
1
0.8
0.6
0.4
0.2
0
pbmc_pDC
pDC
pbmc_Megakaryocyte
pbmc_Monocyte
Monocytes
pbmc_B−cell
Plasmablasts
Memory_B
Naive_B
Naive_Tregs
pbmc_NK_cell
Immature_NK
Mature_NK
Memory_NK
Effector_memory_CD8_T
Gamma_delta_T
pbmc_CD4_T−cell
pbmc_CD8_T−cell
Naive_CD8_T
Naive_Teffs
Central_memory_CD8_T
Th17_precursors
Memory_Teffs
Th1_precursors
Follicular_T_Helper
Th2_precursors
Memory_Tregs
Regulatory_T
PBMC snATAC clustering
FACS sorting (Calderon et al. 2019)
B
Spearman correlation
1
0.8
0.6
0.4
0.2
0
Monocytes
pbmc_Conventional_Dendritic_cell
pbmc_Classical_Monocyte
pbmc_Non−Classical_Monocyte
pbmc_Megakaryocyte
pbmc_pDC
pDC
Plasmablasts
pbmc_Memory_B−cell
pbmc_Naive_B−cell
Memory_B
Naive_B
Effector_memory_CD8_T
Gamma_delta_T
pbmc_Effector_CD8_T−cell
pbmc_Memory_CD8_T−cell
Naive_CD8_T
Naive_Teffs
pbmc_Activated_CD4_T−cell
pbmc_Naive_CD4_T−cell
Central_memory_CD8_T
Th17_precursors
Memory_Teffs
Th1_precursors
Follicular_T_Helper
Th2_precursors
Memory_Tregs
Regulatory_T
Immature_NK
Mature_NK
Memory_NK
pbmc_Adaptive_NK_cell
pbmc_Cytotoxic_NK_cell
pbmc_Regulatory_CD4_T−cell
Naive_Tregs
PBMC snATAC clustering
FACS sorting (Calderon et al. 2019)

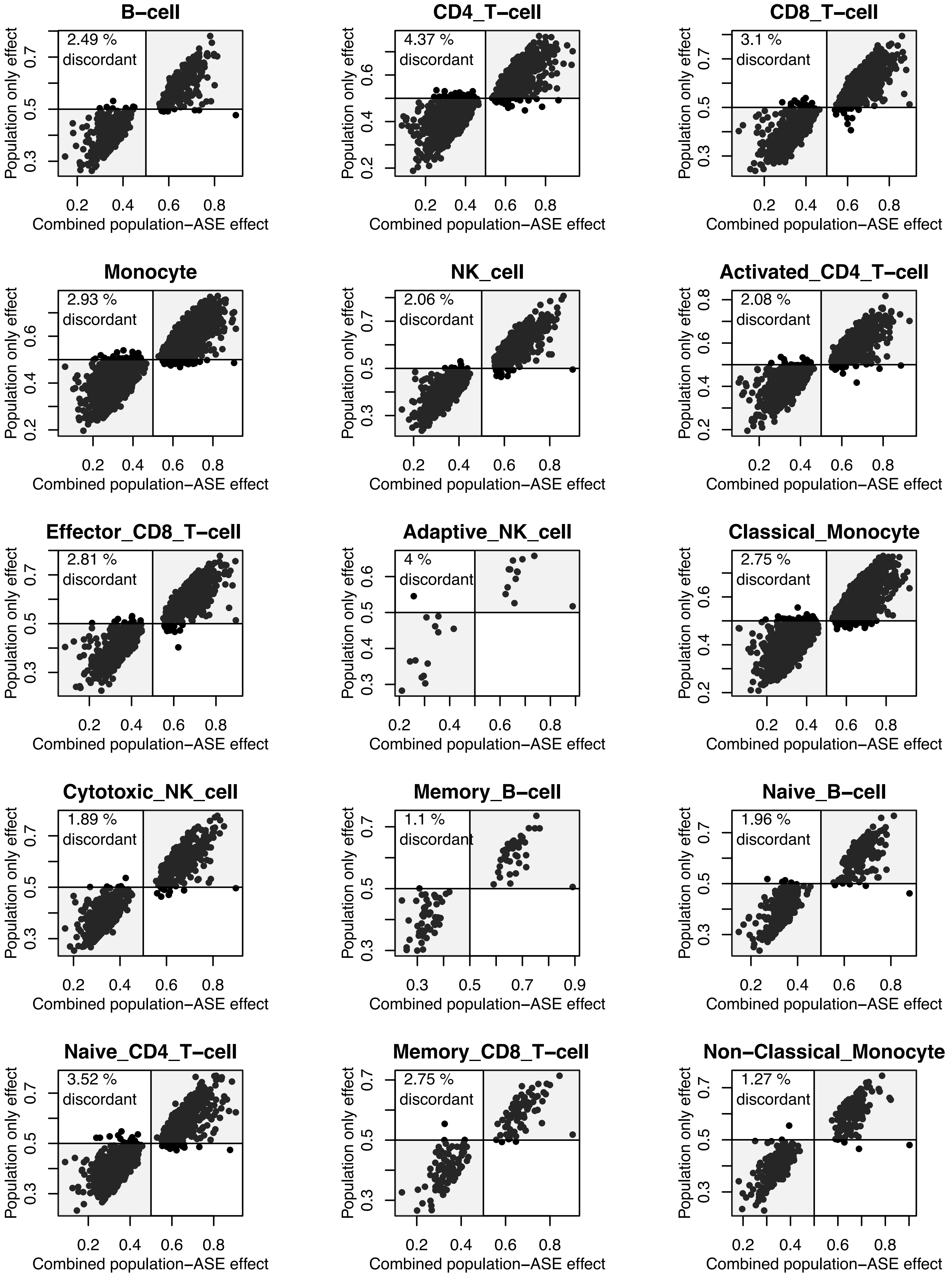
B−cell
2.49 %
discordant
CD4_T−cell
4.37 %
discordant
CD8_T−cell
3.1 %
discordant
Monocyte
2.93 %
discordant
NK_cell
2.06 %
discordant
Activated_CD4_T−cell
2.08 %
discordant
Effector_CD8_T−cell
2.81 %
discordant
Adaptive_NK_cell
4 %
discordant
Classical_Monocyte
2.75 %
discordant
Cytotoxic_NK_cell
1.89 %
discordant
Memory_B−cell
1.1 %
discordant
Naive_B−cell
1.96 %
discordant
Naive_CD4_T−cell
3.52 %
discordant
Memory_CD8_T−cell
2.75 %
discordant
Non−Classical_Monocyte
1.27 %
discordant
Population only effect
Combined population−ASE effect

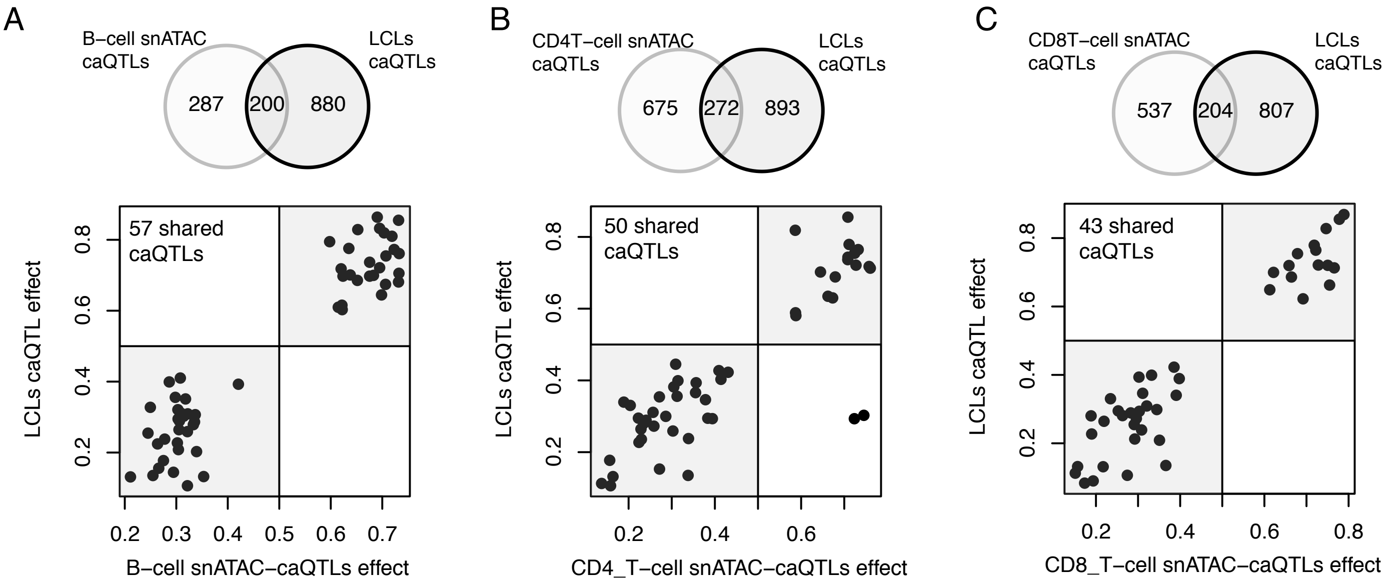


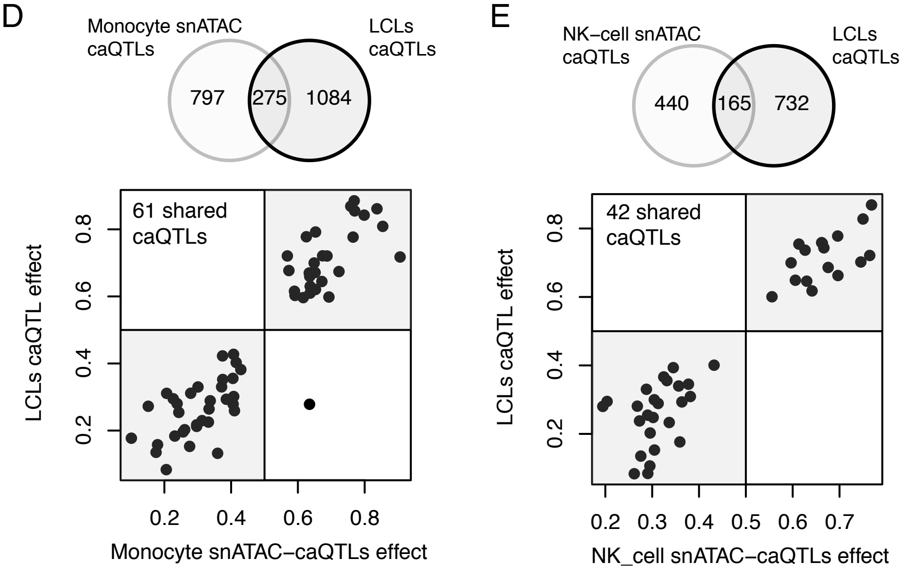


F

Fisher's exact test enrichemnt
of cell-type caQTLs for LCLs caQTLs

| | **P-value** | **Odds Ratio** |
|---|---|---|
| **B-cell** | 5.42E-169 | 25.45809 |
| **CD4_T-cell** | 2.35E-195 | 17.33624 |
| **CD8_T-cell** | 3.46E-142 | 15.85213 |
| **Monocyte** | 1.49E-180 | 14.19069 |
| **NK_cell** | 2.48E-114 | 15.39372 |

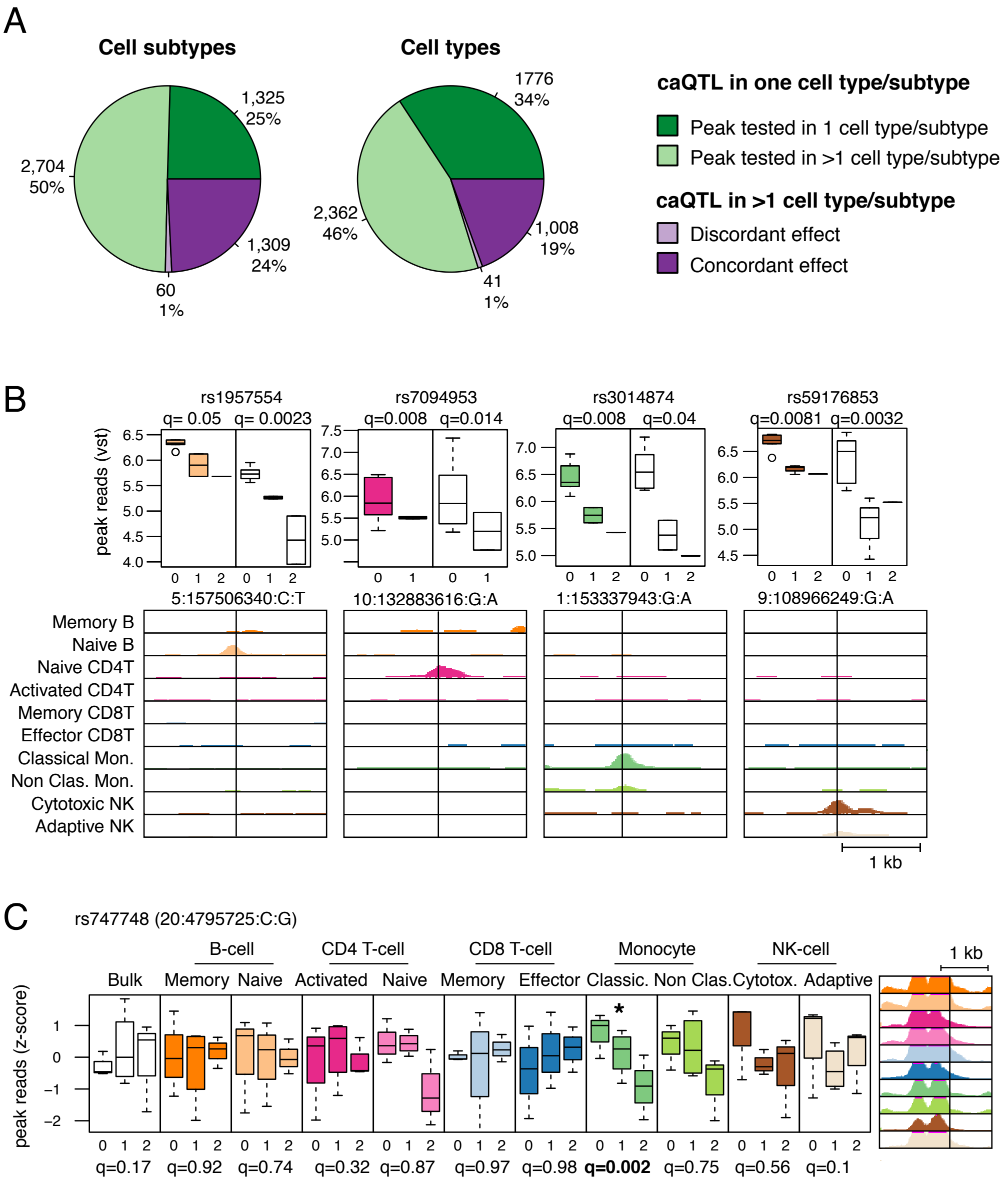

A
Cell subtypes
1,325
25%
2,704
50%
1,309
24%
60
1%
Cell types
1776
34%
2,362
46%
1,008
19%
41
1%
caQTL in one cell type/subtype
Peak tested in 1 cell type/subtype
Peak tested in >1 cell type/subtype
caQTL in >1 cell type/subtype
Discordant effect
Concordant effect
B
rs1957554
q= 0.05 q= 0.0023
rs7094953
q=0.008 q=0.014
rs3014874
q=0.008 q=0.04
rs59176853
q=0.0081 q=0.0032
peak reads (vst)
5:157506340:C:T
10:132883616:G:A
1:153337943:G:A
9:108966249:G:A
Memory B
Naive B
Naive CD4T
Activated CD4T
Memory CD8T
Effector CD8T
Classical Mon.
Non Clas. Mon.
Cytotoxic NK
Adaptive NK
1 kb
C
rs747748 (20:4795725:C:G)
B-cell
CD4 T-cell
CD8 T-cell
Monocyte
NK-cell
Bulk
Memory
Naive
Activated
Naive
Memory
Effector
Classic.
Non Clas.
Cytotox.
Adaptive
1 kb
peak reads (z-score)
q=0.17
q=0.92
q=0.74
q=0.32
q=0.87
q=0.97
q=0.98
q=0.002
q=0.75
q=0.56
q=0.1

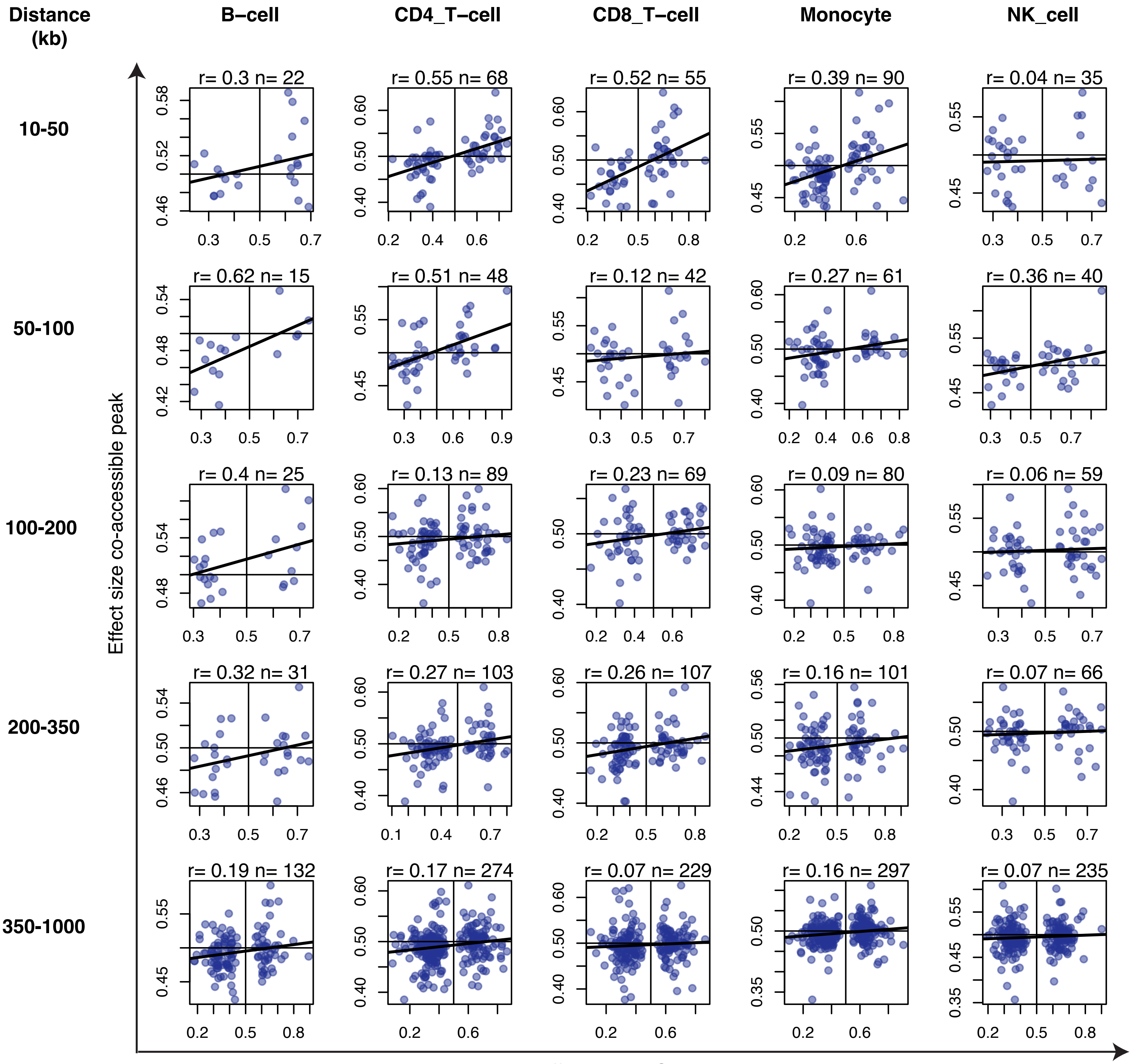

Distance (kb)
B−cell
CD4_T−cell
CD8_T−cell
Monocyte
NK_cell
10-50
r= 0.3 n= 22
r= 0.55 n= 68
r= 0.52 n= 55
r= 0.39 n= 90
r= 0.04 n= 35
50-100
r= 0.62 n= 15
r= 0.51 n= 48
r= 0.12 n= 42
r= 0.27 n= 61
r= 0.36 n= 40
100-200
r= 0.4 n= 25
r= 0.13 n= 89
r= 0.23 n= 69
r= 0.09 n= 80
r= 0.06 n= 59
200-350
r= 0.32 n= 31
r= 0.27 n= 103
r= 0.26 n= 107
r= 0.16 n= 101
r= 0.07 n= 66
350-1000
r= 0.19 n= 132
r= 0.17 n= 274
r= 0.07 n= 229
r= 0.16 n= 297
r= 0.07 n= 235
Effect size co-accessible peak
Effect size caQTL peak

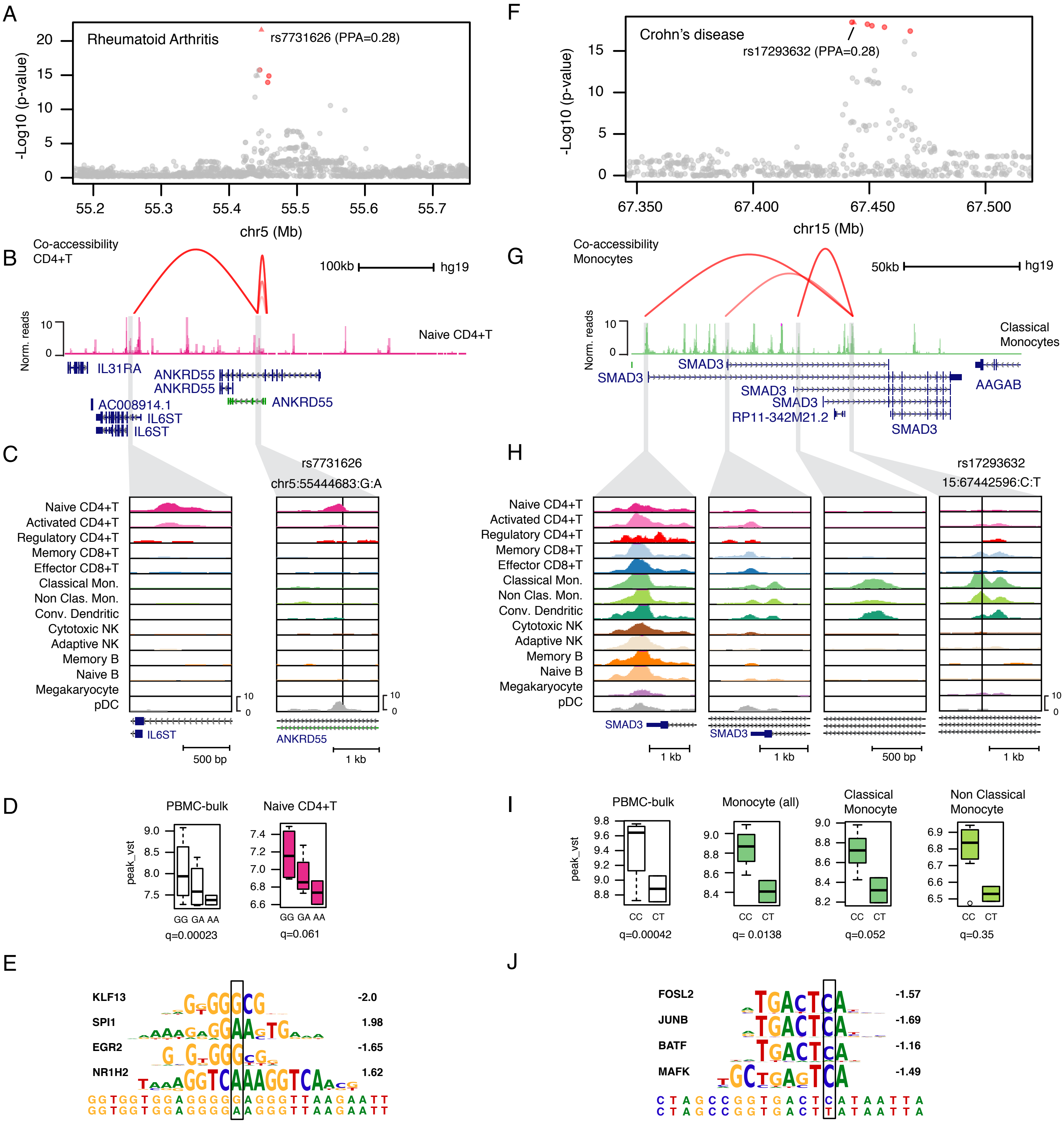
A
Rheumatoid Arthritis
rs7731626 (PPA=0.28)
-Log10 (p-value)
chr5 (Mb)
B
Co-accessibility CD4+T
100kb
hg19
Naive CD4+T
Norm. reads
IL31RA
ANKRD55
AC008914.1
IL6ST
C
rs7731626
chr5:55444683:G:A
Naive CD4+T
Activated CD4+T
Regulatory CD4+T
Memory CD8+T
Effector CD8+T
Classical Mon.
Non Clas. Mon.
Conv. Dendritic
Cytotoxic NK
Adaptive NK
Memory B
Naive B
Megakaryocyte
pDC
500 bp
1 kb
D
PBMC-bulk
Naive CD4+T
peak_vst
GG GA AA
q=0.00023
q=0.061
E
KLF13 -2.0
SPI1 1.98
EGR2 -1.65
NR1H2 1.62
F
Crohn's disease
rs17293632 (PPA=0.28)
chr15 (Mb)
G
Co-accessibility Monocytes
50kb
Classical Monocytes
SMAD3
RP11-342M21.2
AAGAB
H
rs17293632
15:67442596:C:T
I
PBMC-bulk
Monocyte (all)
Classical Monocyte
Non Classical Monocyte
CC CT
q=0.00042
q= 0.0138
q=0.052
q=0.35
J
FOSL2 -1.57
JUNB -1.69
BATF -1.16
MAFK -1.49

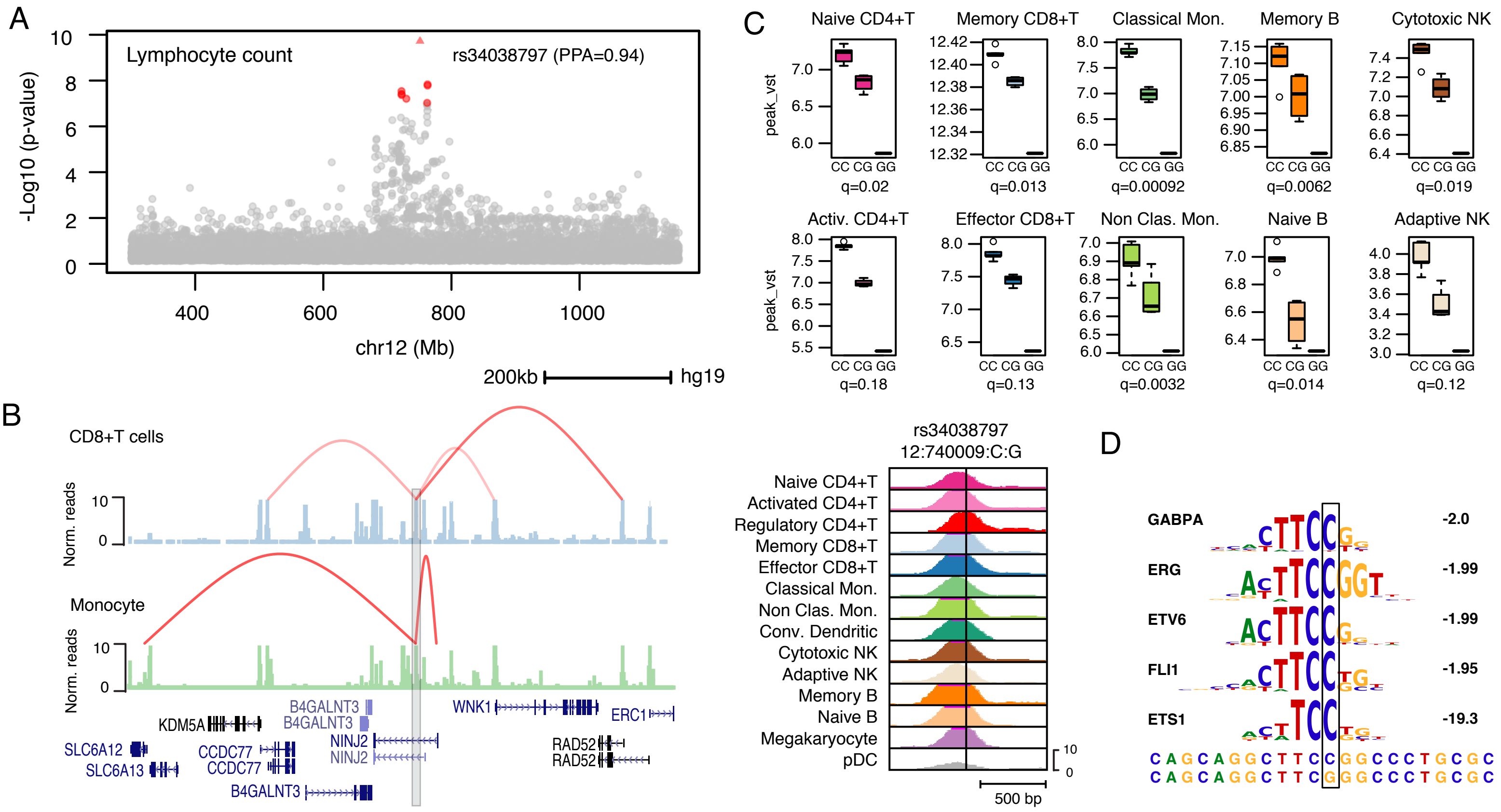

A
Lymphocyte count
rs34038797 (PPA=0.94)
-Log10 (p-value)
chr12 (Mb)
200kb
hg19
B
CD8+T cells
Monocyte
Norm. reads
KDM5A
B4GALNT3
NINJ2
WNK1
ERC1
SLC6A12
SLC6A13
CCDC77
RAD52
C
Naive CD4+T
Memory CD8+T
Classical Mon.
Memory B
Cytotoxic NK
Activ. CD4+T
Effector CD8+T
Non Clas. Mon.
Naive B
Adaptive NK
peak_vst
CC CG GG
q=0.02
q=0.013
q=0.00092
q=0.0062
q=0.019
q=0.18
q=0.13
q=0.0032
q=0.014
q=0.12
rs34038797
12:740009:C:G
Naive CD4+T
Activated CD4+T
Regulatory CD4+T
Memory CD8+T
Effector CD8+T
Classical Mon.
Non Clas. Mon.
Conv. Dendritic
Cytotoxic NK
Adaptive NK
Memory B
Naive B
Megakaryocyte
pDC
500 bp
D
GABPA -2.0
ERG -1.99
ETV6 -1.99
FLI1 -1.95
ETS1 -19.3
CAGCAGGCTTCCGGCCCTGCGC
CAGCAGGCTTCGGGCCCTGCGC
